# An updated review of the marine ornamental fish trade in the European Union

**DOI:** 10.1101/2024.03.17.585413

**Authors:** Monica V. Biondo, Rainer P. Burki, Francisco Aguayo, Ricardo Calado

**Affiliations:** Fondation Franz Weber, Bern, Switzerland; asdfg IT, Fluh 86, Rosshäusern, Switzerland; Faculty of Higher Studies Cuautitlán, National Autonomous University of Mexico, Mexico City; ECOMARE, CESAM—Centre for Environmental and Marine Studies, Department of Biology, Santiago University Campus, University of Aveiro, Aveiro, Portugal

**Keywords:** Coral reef fishes, international aquarium trade, management, value, TRAde Control and Expert System TRACES

## Abstract

Wild-caught fishes from coral reefs, one of the most threatened ecosystems on the planet, continue to supply the marine aquarium trade. Despite customs and veterinary checks during imports, comprehensive data on this global industry remains scarce. This study provides an updated review on one of its largest import markets, the European Union (EU): 24-million-euro annual trade value, detailed exporting and importing countries also diversity of species and families of the 26 million specimens imported between 2014 and 2021. We then show how a watchlist alert system based on the number of specimens traded, import trends, and vulnerability index according to FishBase and IUCN Red List conservation status can provide key information on which species should require a closer scrutiny. The stark decline in trade of the most traded species, *Chromis viridis,* may warrant monitoring through the Convention on International Trade in Endangered Species of Wild Fauna and Flora (CITES). While the European TRAde Control and Expert System (TRACES) electronically monitors the movement of live animals to respond quickly to biosecurity risks, one-third of marine ornamental fishes imported lack species-level information. With minor adjustments, TRACES could enhance the monitoring of wildlife trade, with marine ornamental fish being an interesting case study.

## Introduction

The European Union (EU) is a major player in the global market for marine ornamental fishes, both in terms of value and number of specimens. This trade has been expanding (Bruckner 2005; Thornhill 2012; Kültz 2022) and leading to further pressure on species and ecosystems. The EU Parliament’s resolution of 5 October 2022 emphasized the importance of addressing the trade of marine ornamental fishes and its monitoring, particularly as most of marine ornamental fishes are wild caught (EU 2022a). Additionally, during the 18^th^ Conference of the Parties to the Convention on International Trade of Endangered Species of Wild Fauna and Flora (CITES) in August 2019, it was recognized that the international trade of marine ornamental fishes required closer examination due to its large scale, lack of regulation, and inadequate monitoring (CITES 2019). A workshop involving CITES parties, industry representatives, experts, and NGOs was planned, but its conclusion was delayed until the 20th CITES conference in 2025 due to the COVID-19 pandemic (CITES 2021).

Since 2004, the EU has electronically monitored the movement of live animals, plants, and food from third party countries (non-EU countries) to the EU using the TRAde Control and Expert System (TRACES). This system enables the EU to respond quickly to biosecurity risks like potential threats posed by zoonosis and invasive species (TRACES 2023). In 2014, the monitoring of marine ornamental fishes through TRACES became possible with the introduction of the “Harmonized System code 03011900 for Live ornamental fish (excluding freshwater)”. However, from 2014 to 2021 one third of imported fishes lacked species-level information, making it impossible to assess any biosecurity risks associated with these imports (Biondo and Burki 2019; Biondo and Calado 2021). Nonetheless, this data collection system provides an overview of this trade into Europe despite some shortcomings.

All fishes imported into the EU are transported by air freight, primarily from southeast Asia (Indonesia, the Philippines, and Sri Lanka). Upon arrival in the EU, these specimens undergo customs clearance, as well as veterinary inspection, and must be registered in the European TRAde Control and Expert System (TRACES) before being collected by wholesalers or buyers (Biondo and Burki 2019; Biondo and Calado 2021).

The global trade of marine ornamental fishes, spanning nearly a century, has never been effectively monitored (Biondo 2017, 2018; Biondo and Burki 2019, 2020; Dee et al. 2014; Rhyne et al. 2012, 2017; Wabnitz et al. 2003), as previous attempts to increase transparency and oversight within the industry have been unsuccessful (Murray et al. 2012; Townsend 2011; UNEP-WCMC 2008). This complex industry involves over 60 export countries and targets more than 2,000 species (Rhyne et al. 2012), but the environmental impacts of harvesting millions of coral reef fishes per year for this trade have largely been overlooked (Grutter et al. 2018; Thornhill 2012; Tissot et al. 2010; UNEP-WCMC 2008; Vagelli 2011). Some species have already been negatively impacted by the marine aquarium trade (Hinsley et al. 2023). For instance, the Banggai cardinalfish (*Pterapogon kauderni*) was discovered by the aquarium industry in the 1990s and has come perilously close to extinction due to its high demand (Vagelli 2011). In consequence, it was listed as “endangered” on the IUCN Red List in 2007 and the United States had placed it on its Endangered Species Act and listed it as “threatened” in 2016 (NOAA 2016); presently the US are calling for additional protection by proposing to ban imports and exports of wild and captive-bred specimens of *P. kauderni* (NOAA 2023). Another documented example of the impact of this trade on some species is the overfishing of bluestreak cleaner wrasse (*Labroides dimidiatus)* and its fostering of biodiversity loss on coral reefs (Bshary 2003; Grutter et al. 2018; Waldie et al. 2011). Ongoing climate change threatens coral reefs and the marine ornamental fish species they host (Carpenter et al. 2008; Ferrari et al. 2011; Gattuso et al. 2014; Hoegh-Guldberg et al. 2019; Hughes et al. 2017; IPCC 2018; Souter et al. 2020). The impacts of destructive fishing practices used in the marine aquarium industry, such as cyanide fishing, are further exacerbated by warmer waters, as suggested by some laboratory studies (Madeira et al. 2020). High mortality rates along the supply chain also contribute to the decline of marine ornamental fish populations, as extra specimens need to be collected to cover such losses (Cohen et al. 2013; Conant 2015; Dee et al. 2014; Huntingford et al. 2006; Monticini 2010; Vagelli, 2011; Wabnitz et al. 2003).

In this study, we analyzed EuroStat and UN Comtrade data to assess the monetary value of the global trade in marine ornamental fishes, the importance of the EU demand in this trade and the key trends of this market which predominantly relies on wild-caught specimens. Additionally, we utilized TRACES data from 2014 to 2017 (Biondo and Burki 2019) and analyzed new data from 2018 to 2021, retrieving information available on country of origin and destination (export and import country), species diversity, number of specimens, and trade trends. By considering the vulnerability of species based on FishBase, and their conservation status according to the IUCN Red List, we developed an alarm system, a three parameters list and called it Watchlist: including a previous study covering data from 2014 to 2017 (Biondo and Burki 2019). Finally, we extend this to a Watchlist+ (including a linear regression for estimating the time-trend in number of specimens traded) aimed to rank species that may be at risk of overexploitation due to the global marine aquarium trade’s impact on Europe’s imports. Our goal is to shed light on this century old trade with a focus on Europe, using the best available data with regards to value, number of specimens, and diversity and show a way forward to reliably monitor this global trade.

## Materials and methods

### Data on marine ornamental fish value

To assess the economic value of trade in marine ornamental fishes, we examined first the import values at the global and regional level using the nominal value of imports (in US$) from the United Nations Commodity Trade Statistics Database (UN-Comtrade database). This source is compiled by the United Nations Statistics Division from detailed global annual and monthly trade statistics by product and trading partner, covering approximately 195 countries, and representing more than 99% of the world’s merchandise trade (UN 2023). We extracted import values for the product classification “Harmonized System code 03011900 Live ornamental fish (excl. freshwater)” (HS 03011900) for the period 2014 to 2021. Global imports reported by country were aggregated by region based on the World Bank’s regional classification (World Bank 2023). China was separated from the rest of East Asia and the Pacific because of its size and for showing a distinctly different (rapidly increasing) trend with respect to the region. For the EU region, we included the 27 EU countries and the UK (up to the end of 2020, as beyond this date this country was no longer an EU member-state), as well as Iceland, Norway, San Marino, and Switzerland. We converted import values into euros (€) using annual average exchanges rate of the US$.

It must be noted that import values from UN-Comtrade have known biases, there is considerable under-reporting and import values are most likely over-estimated as many products are re-exported (Chan et al. 2015; Andersson et al. 2021). Correcting this bias directly was not possible since re-exports are practically un-reported in this database. This is an important source of bias given the high level of trade within the EU (imports from the rest of the world that are re-exported to another EU member state). As described with detail in Leal et al. (2015), the structure of imports within the EU reflects a high degree of specialization in the trade of marine ornamental fishes and allows to clearly distinguish between exporter (UK, Netherlands, Germany) and importer countries (Spain, Italy). To avoid re-exports from being double counted as imports, we approximate the value of extra-EU imports by taking the percentage they represent in total EU imports from the EuroStat database (as reported in the Statistical Office of the European Communities’ database EU trade since 1988 by HS2-4-6 and CN8 (ds-045409), EUROSTAT 2023). We estimate the value of extra-EU imports by applying this percentage to the value of total EU imports as reported in UN-Comtrade.^1^

Import values are only a fraction of the market size. After being imported, value is added to the product (in this case, the fishes) as wholesalers and retail traders add their costs to the selling price. This added value can be approximated in turn by calculating the net exports, the difference between EU exports and imports as reported in UN-Comtrade. The trade value chain ends with final consumers; this final demand is a better approximation to the size of the market. To approximate the value of the final demand for ornamental marine fishes in the EU we take the value of extra EU-imports as described above and add the value of net-exports. The latter can be interpreted as a measure of the costs added by traders to the final price. To calculate average prices in Euros we used the extra-EU import value and the total import value from Eurostat. All figures in euros are discounted for inflation using the Harmonized Index of Consumer Prices (EuroStat, 2023 dataset: HICP – annual data, average index, and rate of change) set to 2020 = 100, so that all values are comparable over time.

### Data on marine ornamental fish species and numbers

As for data from 2014-2017 (Biondo and Burki 2019), the EU Directorate-General for Health and Food Safety (DG SANTE) provided Excel files containing data on marine ornamental fishes imported to Europe from 2018 to 2021, obtained from the European TRAde Control and Expert System TRACES using “HS 03011900 Live ornamental fish (excluding freshwater)”. TRACES is an online platform used for sanitary certification and traceability of imports of live animals, animal products, food, and feed into Europe. It helps mitigate biosecurity risks and disease outbreaks (TRACES 2023). Introduced in 2004, TRACES is widely utilized in approximately 90 countries by 55,000 users (government agencies, exporting and importing businesses, official veterinarians) and is available in 39 languages. It facilitates cooperation between EU and non-EU authorities. Since 2014, TRACES has been gathering data specifically for marine ornamental fishes under the category “HS 03011900 Live ornamental fish (excluding freshwater)”, which was previously grouped more broadly as “otra pesca” (Biondo 2017).

Traders are required to be registered with TRACES and complete customs documents which also physically accompany consignments. The import goods are declared at the border of any EU member state, as well as Switzerland, Norway, Iceland, or San Marino, by entering the freight details in the web interface of TRACES. TRACES is the only tool to collect information on number of specimens and diversity of marine ornamental fishes in the European region, although without securing a true traceability for these marine organisms. Until the end of 2019 customs documents were titled as “Common Veterinary Entry Document Animals” (CVEDA) and then became the harmonized “Common Health Entry Document Animals” (CHEDA), designed specifically to carry out health checks at borders. This approach led to an adaptation of TRACES data being collected for marine ornamental fishes with regards to data for 2014 to 2017, allowing the import of Excel files and making possible the use of taxonomic higher-level data, including at the order level.

The data provided by the EU containing freshwater ornamental fishes, invertebrates, and other non-fishes, along with rejected shipments, were excluded from this study. Species from land-locked countries or countries with no tropical waters were retained, as these countries could represent commercial transit hubs for this trade. The record signaling 80,000 *Muraena helena* shipped from Israel to Denmark were removed from the watchlists as these were not clearly destined for aquaria (*M. helena* is a regular food fish; because of its size only a few are kept in public aquariums). One record of a shipment destined for the United Kingdom but having as its destination country the United States of America, was changed to the United Kingdom, as the border inspection post was in the UK. Countries with multiple possible names were harmonized: “United Kingdom (Northern Ireland)” and “The Netherlands” were changed to “United Kingdom” and “Netherlands”, respectively.

All scientific fish names were checked by using the World Register of Marine Species (WoRMS; http://www.marinespecies.org) and FishBase (http://www.fishbase.org) the global species database of fish species. Records of fishes lacking their complete genus or species identification were allocated to their family, as well as all fishes with complete species names, by using FishBase and WoRMS.

TRACES data were cleaned by using information from FishBase by filtering out fishes that did not match “saltwater” AND (“tropical” OR “subtropical”) AND “reef-associated” plus “saltwater” AND (“tropical” OR “subtropical”) AND “demersal” AND “aquarium”. Cichlidae or Toxotidae are primarily freshwater fishes but FishBase either places the species as “reef-associated” (brackish) and/or as migratory ocean-river. Scophthalmidae are commonly traded as food fishes, but these specimens were imported as ornamentals and were therefore considered as well in the present study. Information on origin and destination, number of specimens traded, species diversity were analyzed.

### Trends in number of specimens traded and watchlists

Two separate lists were produced: a Watchlist and a Watchlist+. For the Watchlist, the score for each species was evaluated through three parameters: number of specimens imported per year, vulnerability according to FishBase, and the IUCN Red List conservation status (Biondo and Burki 2019). The median number of specimens traded was normalized, assigning a value of 100 to the species with the highest four-year median trade volume. Data outliers were retained as they may be accurate. The IUCN Red List categories were converted into numerical values as follows: “least concern” (LC) = 0, “near threatened” (NT) = 20, “vulnerable” (VU) = 40, “endangered” (EN) = 60, “critically endangered” (CR) = 80, and “extinct in the wild” (EW) = 100. “Extinct” (EX) was not assigned a value since trading an extinct species is not possible. For “data deficient” (DD) or “not evaluated” (NE) species the IUCN preamble states: “*until such time as an assessment is made, taxa listed in these categories should not be treated as if they were non-threatened. It may be appropriate (especially for* “*data deficient*” *forms) to give those species the same degree of attention as threatened taxa, at least until their status can be assessed.*” For this and as the habitat of marine ornamental fishes, coral reefs, are threatened (IPCC 2018; Souter et al. 2020) these categories were handled as “vulnerable” (VU) and received the numerical value of 40. FishBase computes a vulnerability score for each species, representing its resilience to external factors and is calculated using selected life-history parameters, with a score ranging from 0 to 100. A score was determined for each species by summing the normalized parameter values. The higher the score the more potentially exposed the species could be to overexploitation through the fishing effort to supply the marine aquarium trade. As every category is normalized to a value between 0 and 100, the theoretical maximum would be a score of 300. By 2022 the IUCN Red List had re-evaluated 449 species, with the following results: 1 species was newly rated as “endangered” (EN), 4 as “vulnerable” (VU), 2 as “near threatened” (NT), 426 as “least concern” (LC) and 16 as “data deficient” (DD). This information was updated in the data spreadsheet prior to analysis.

The Watchlist+ was produced using the same three parameters detailed above for the Watchlist but including a linear estimate of the trend in the number of specimens traded over the eight years (slope). The linear regression was tested for significance with a weighted R-squared, for variance explanatory power and a t-test on the coefficient significance. Only species with a p-value for the t-statistic of less or equal 0.05 were retained. The time series has only 8 observations, which may be just enough for ensuring statistical significance if variance is low (Jenkins and Quintana-Ascencio, 2020). The Watchlist+ revealed that for several species of marine ornamental fishes slope estimations were not meaningful due to high variance or missing data. This method will however improve in accuracy as the database grows in time.

## Results

### Import values of marine ornamental fishes

According to UN-Comtrade, between 2014 and 2021 the EU total import values of marine ornamental fishes reached €24 million in average per year (at 2020 constant prices). This value includes the UK (up to the end of 2020), as well as Iceland, Norway, San Marino, and Switzerland (for the whole period 2014-2021). If we consider imports coming strictly from outside the EU-27 (“extra-EU”, Figure 1), the figure is reduced to €12.1 million in average between 2014-2021. Although its share in the global imports of these organisms has diminished (from 40.5% in 2014) the EU still accounts for 35.4% of the global imports in 2021 (Figure 2). As shown in Figure 3, EU import values have a cyclical trajectory with an almost constant trend, while trade in the rest of the world exhibits a clear increasing trend, driven mostly by North America and China. The value of extra-EU imports decreased by 26.8% between 2014 and 2021 (Figure 4), which is aligned with the 59.9% decrease in the number of specimens reported (Figure 5). Final demand, however, increased by 14.6% during the same period, averaging €35.4 million in 2014-2021 (Figure 4), and does not exhibit a reducing trend during the whole period 2014-2021. Based on the observed data for value and number of specimens, average import prices doubled from €2.1 to €4.6 per specimen between 2014 and 2021, when considering only extra-EU imports, but more than doubled from €6.9 to €19.8 per specimen when final demand is considered (Figure 6).

**FIGURE 1.**
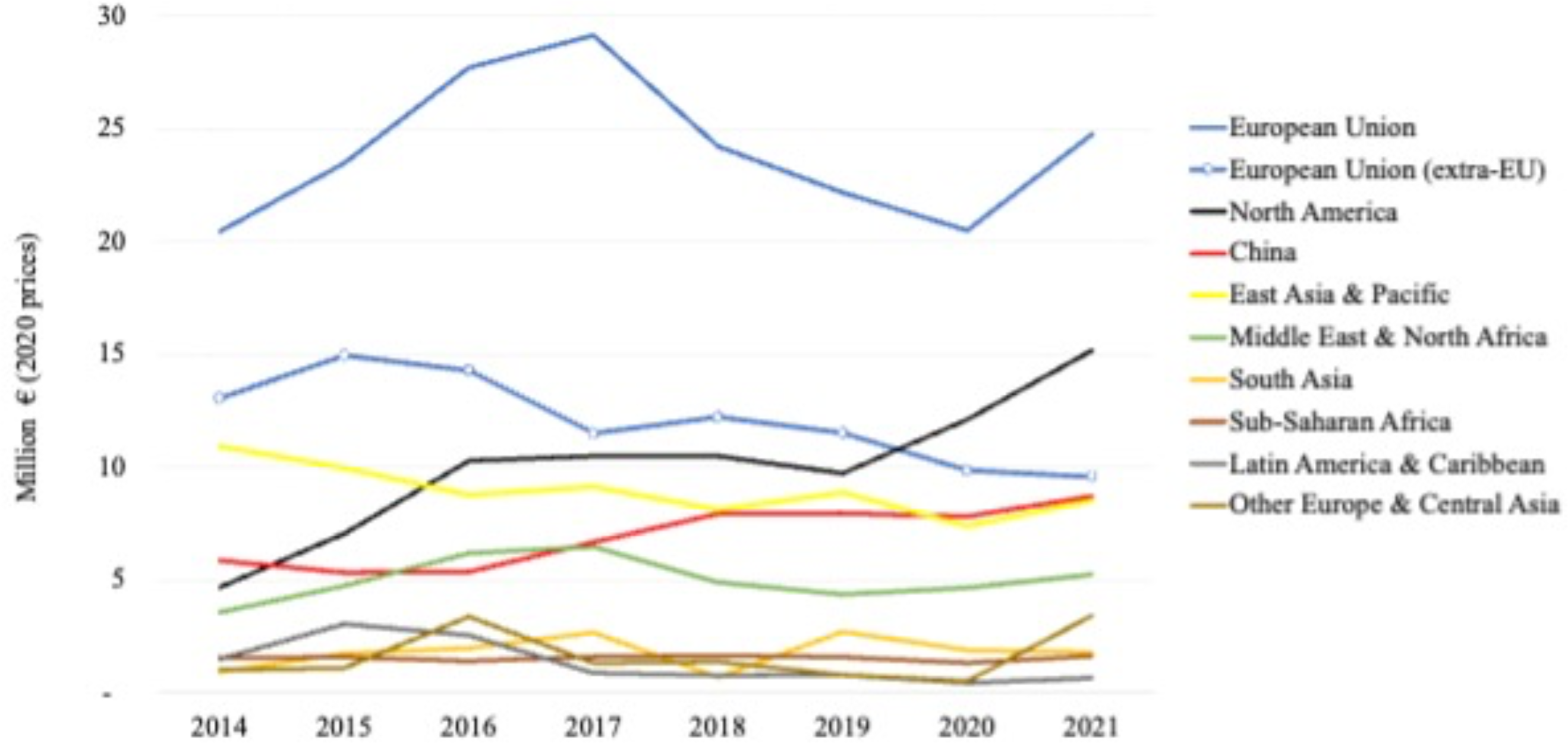
Global imports by region 2014-2021 in million € (2020 prices). Notes: 1) European Union includes: 27 EU countries plus UK (until 2020), Switzerland, Norway, Iceland and San Marino (UN-Comtrade database, code “HS 030119: Fish; live, ornamental, other than freshwater”). 2) Import values are deflated by the Harmonized Index of Consumer Prices (EuroStat 2023).

**FIGURE 2.**
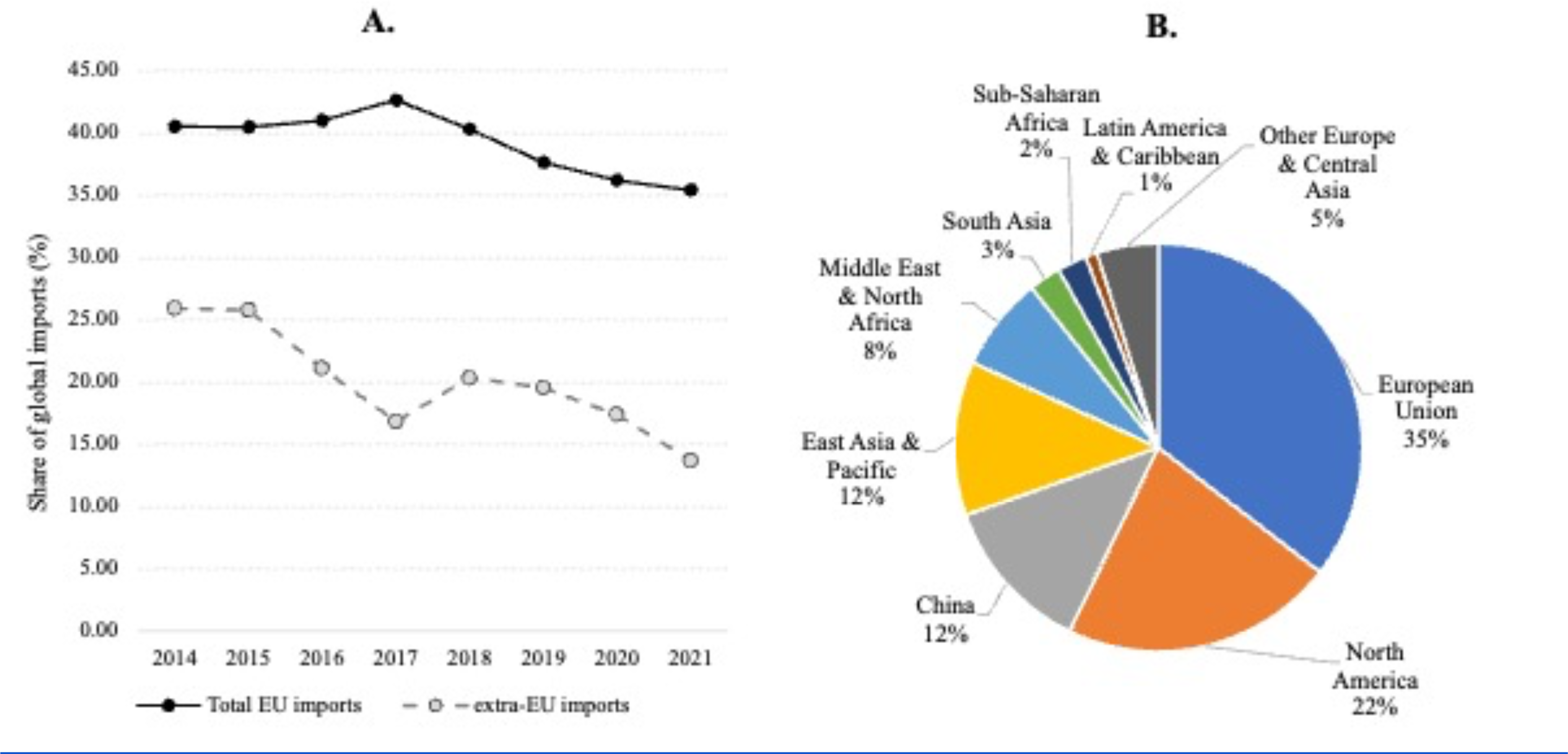
A: Market share of European imports in the global imports value of marine ornamental fishes. B: Share (%) of import value by region in 2021. Source: own calculations based on UN-Comtrade database, “HS classification 030119: Fish; live, ornamental, other than freshwater”.

**FIGURE 3.**
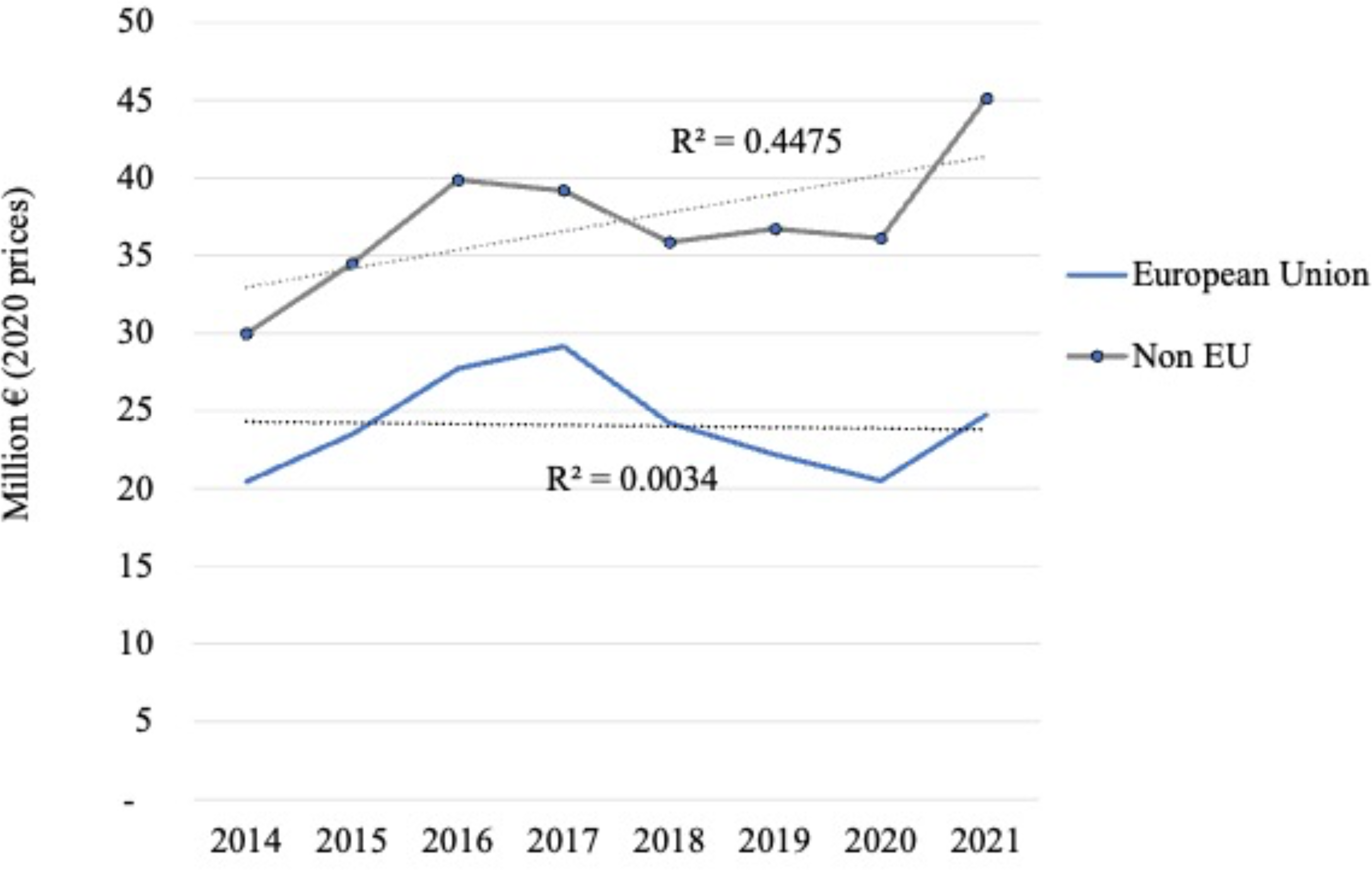
Global trade and linear regression (dotted line) in marine ornamental fishes: Annual imports by region 2014-2021 (million € at 2020 prices); own calculations based on UN-Comtrade database, classification “HS 030119: Fish; live, ornamental, other than freshwater”.

**FIGURE 4.**
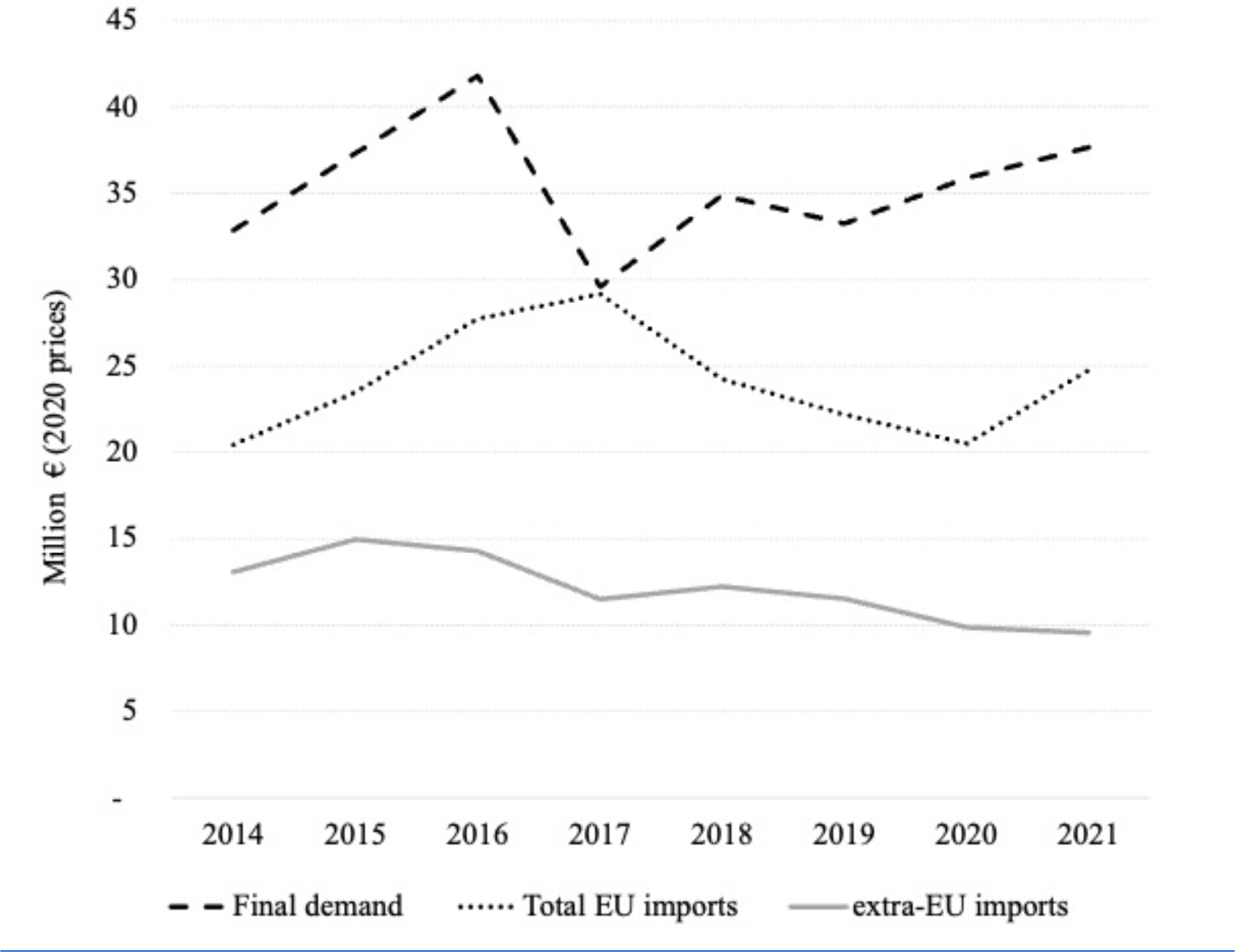
European value trade (imports) and final demand in marine ornamental fishes (million € at 2020 prices) from 2014 to 2021; authors’ calculations based on UN-Comtrade database, classification “HS 030119: Fish; live, ornamental, other than freshwater”.

**FIGURE 5.**
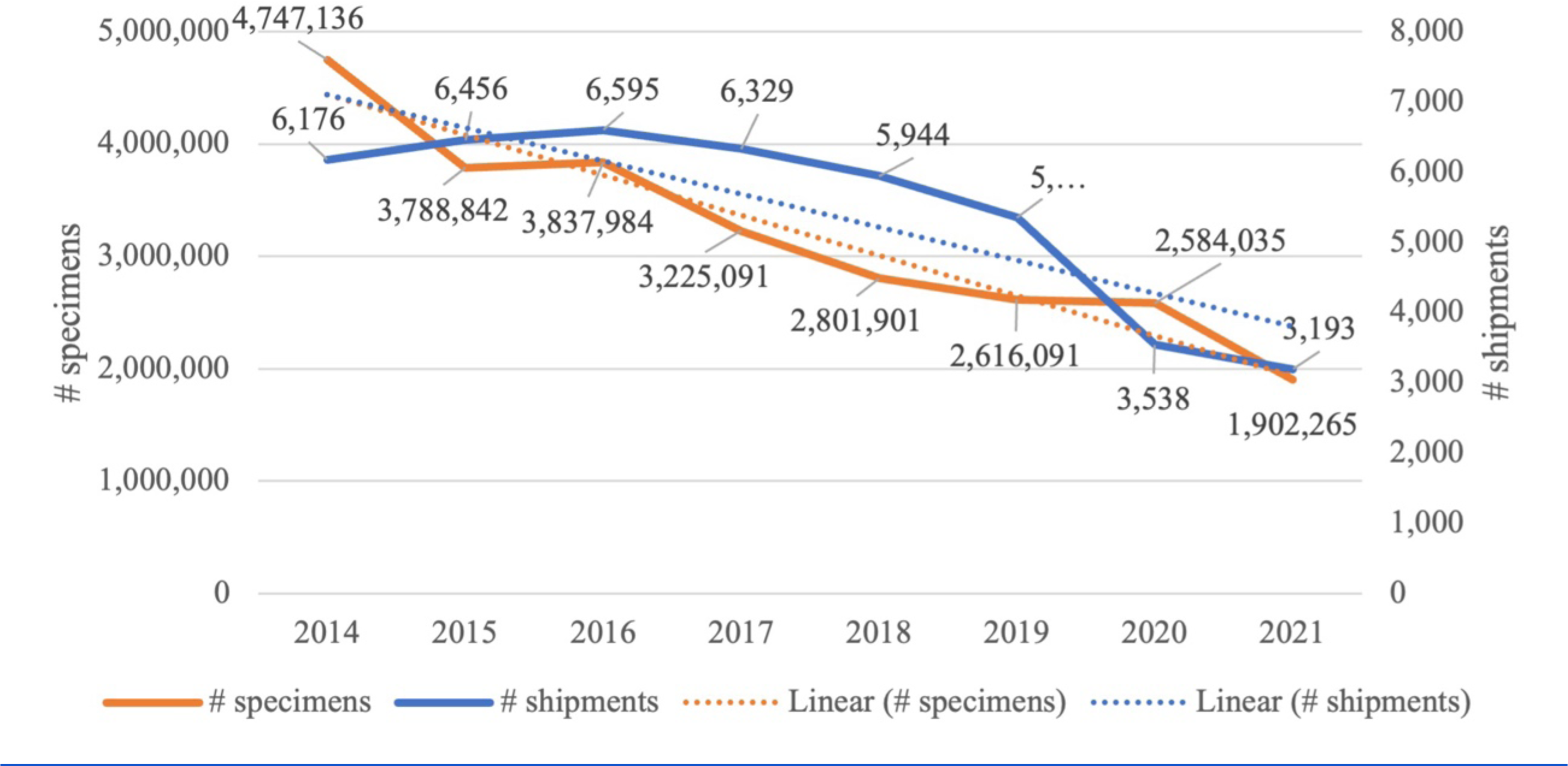
Overall shipments and number of traded specimens per year of marine ornamental fishes and linear regression from 2014 to 2021 entering Europe according to TRACES data.

**FIGURE 6.**
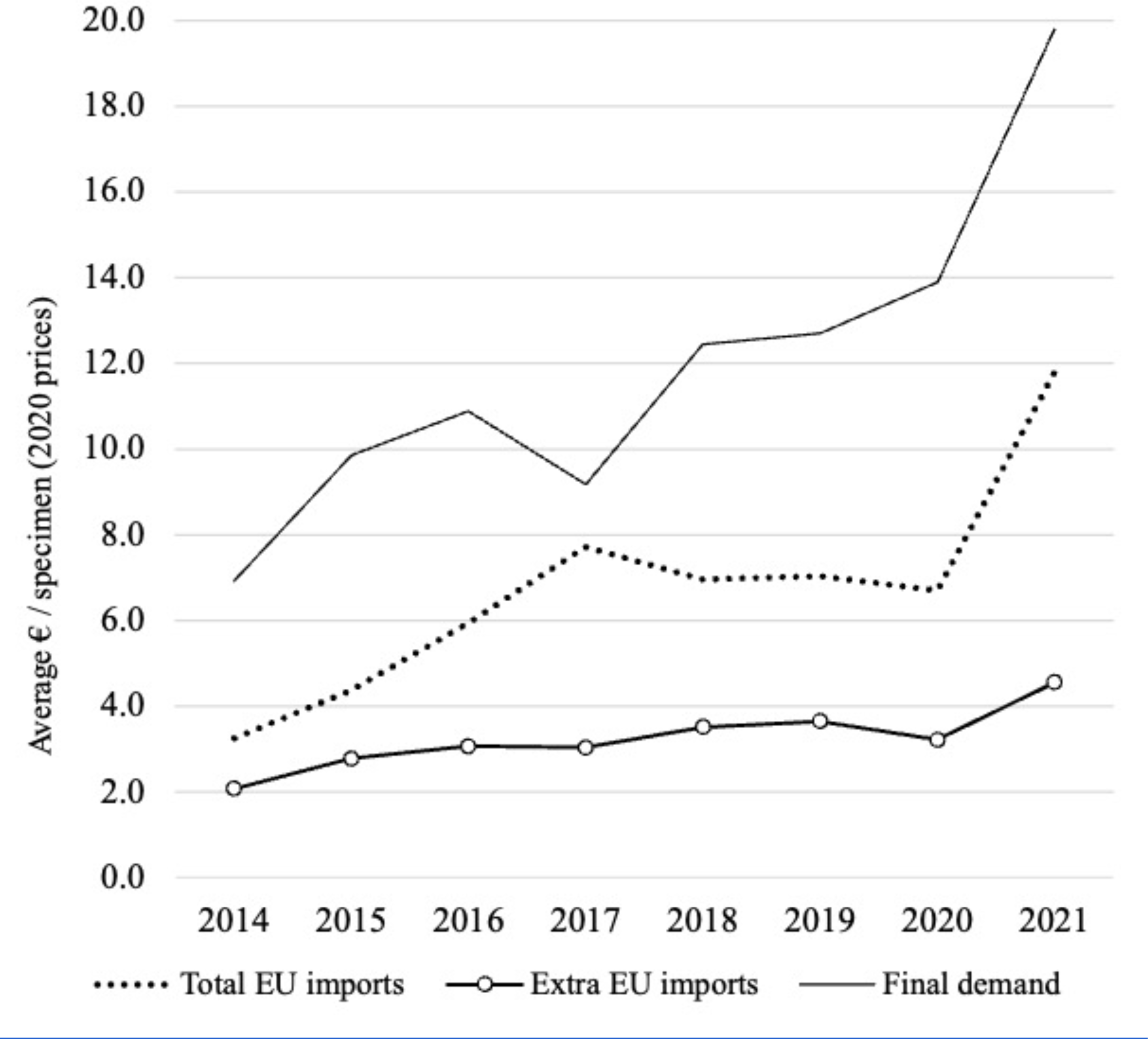
European marine ornamental imports: average prices and final demand in marine ornamental fishes (value in €, 2020 prices) from 2014 to 2021; authors’ calculations based on Eurostat database and TRACES data.

### Import data according to TRACES

The analysis of TRACES data from 2014 to 2021 revealed a total of 297,412 data records; a record being a shipment item consisting of a species and the number of fishes shipped (Table 1). A shipment often included multiple species. A total of 282,226 records represented 25,503,345 marine ornamental fishes, whilst 15,186 were not of marine origin and included freshwater ornamental fishes, invertebrates, with even 5 records of amphibia (Table 1). Overall, 59.9% of all ornamental fish specimens registered were indeed marine ornamental fishes (Table 1).

**TABLE 1.**
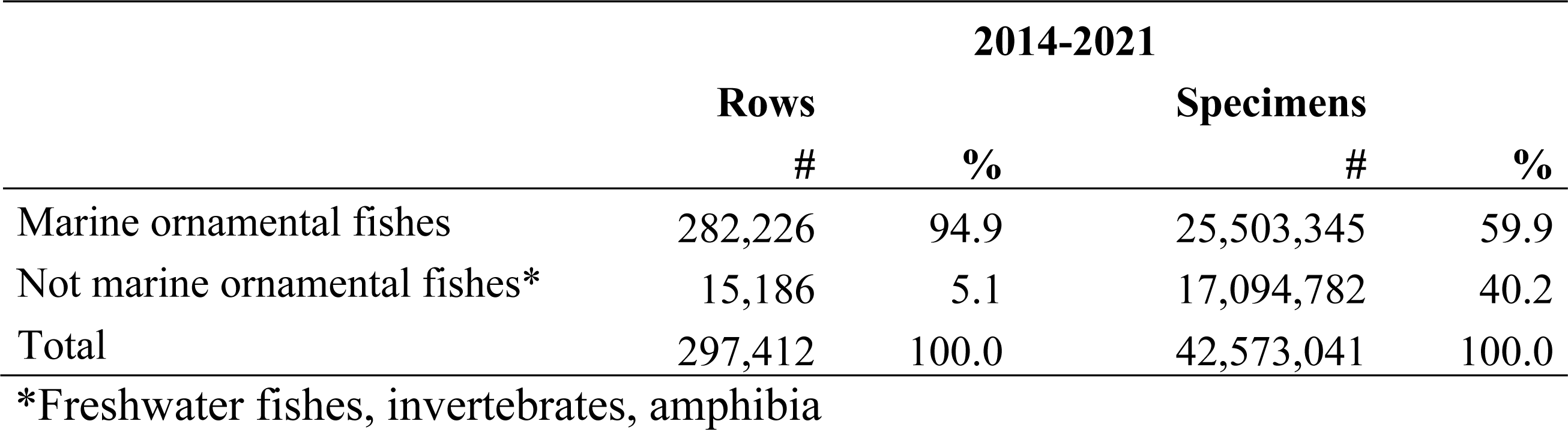
Units (rows) of imports into the EU of specimens of marine and non-marine ornamental fishes, invertebrates, and amphibians for 2014 to 2021 according to TRACES data.

|  | 2014-2021 |  |  |  |
| --- | --- | --- | --- | --- |
|  | Rows |  | Specimens |  |
|  | # | % | # | % |
| Marine ornamental fishes | 282,226 | 94.9 | 25,503,345 | 59.9 |
| Not marine ornamental fishes* | 15,186 | 5.1 | 17,094,782 | 40.2 |
| Total | 297,412 | 100.0 | 42,573,041 | 100.0 |

\*Freshwater fishes, invertebrates, amphibia

### Country of origin, destination, and specimens

From 2014 to 2021 a total of 61 countries exported marine ornamental fishes to Europe. The main exporting country was Indonesia with 33.5% of shipments and an average of 1,394,208 specimens exported per year (Table 2). With regards to the number of specimens exported, Indonesia was followed by the Philippines with an average of 529,076 fishes and Sri Lanka with 266,945 fishes; however, in terms of shipments, Sri Lanka displayed a higher number than the Philippines (16.2% and 12.2%, respectively). The same trend was recorded for the Maldives, Singapore, and Israel, with the Maldives exporting higher fish numbers but Singapore and Israel making more shipments.

**TABLE 2.** Top eight exporting countries of marine ornamental fishes to Europe between 2014 and 2021. Average, standard deviation and % of shipments and specimens, and value of imports per year. Total imports refer to imports from outside the EU 27. AVG = average, SD = standard deviation. Value of imports, € (2020 prices). Total = extra EU 27 imports, excluding intra-EU trade; EuroStat.

| Country of origin | Shipments |  |  | Specimens |  |  | Value of imports (€) |  |  |
| --- | --- | --- | --- | --- | --- | --- | --- | --- | --- |
|  | AVG | SD | % | AVG | SD | % | AVG | SD | % |
| Indonesia | 1,823.6 | 436.8 | 33.5 | 1,394,208 | 393,487 | 43.7 | 3,621,793 | 527,884 | 38.1 |
| Philippines | 663.5 | 183.6 | 12.2 | 529,076 | 100,105 | 16.6 | 888,489 | 133,226 | 9.3 |
| Sri Lanka | 883.3 | 175.6 | 16.2 | 266,945 | 62,842 | 8.4 | 863,273 | 80,583 | 9.1 |
| United States | 441.8 | 162.6 | 8.1 | 221,012 | 131,618 | 6.9 | 1,235,212 | 384,862 | 13.0 |
| Kenya | 391.0 | 74.7 | 7.2 | 197,073 | 21,001 | 6.2 | 791,417 | 300,613 | 8.3 |
| Maldives | 198.8 | 66.9 | 3.6 | 118,709 | 26,912 | 3.7 | 234,698 | 50,275 | 2.5 |
| Singapore | 356.8 | 123.4 | 6.5 | 107,222 | 103,373 | 3.4 | 118,618 | 44,016 | 1.3 |
| Israel | 236.6 | 91.8 | 4.3 | 86,151 | 31,104 | 2.7 | 146,374 | 43,452 | 1.5 |
| Fiji | 58.3 | 23.9 | 1.1 | 46,633 | 21,231 | 1.5 | 51,124 | 15,433 | 0.5 |
| Dominican Republic | 57.5 | 28.1 | 1.1 | 37,638 | 31,289 | 1.2 | 26,789 | 18,208 | 0.3 |
| Other | 12.4 |  | 6.2 | 6,756 |  | 5.7 | 1,531,137 |  | 16.1 |
| Total |  |  | 100.0 |  |  | 100.0 |  |  | 100.0 |

The three main exporting countries, Indonesia, the Philippines, and Sri Lanka accounted for 61.9% of shipments and 68.7% of specimens and, together with the United States, Kenya, the the Maldives, Singapore and Israel they summed up 91.7% of all shipments and 91.6% of all specimens of marine ornamental fishes entering Europe (Table 2). These eight countries represented 83.9% of import’ value (Table 2). In terms of the value of trade, the United States came second and the Philippines third in importance after Indonesia, shipping fewer specimens but fetching higher average prices.

With regards to value, on average, Indonesia shipped 765 specimens per shipment at a value of € 2.6 per fish whereas the Philippines shipped more species per shipment, 797 specimens, with each fish costing € 1.7 (Table 3). The most expensive fishes were shipped from the United States at an average value of € 5.6 per fish (Table 3).

**TABLE 3.** Average number of specimens per shipment and average price per specimen in € (2020 prices) of the top eight exporting countries between 2014 and 2021; EuroStat.

| **Country of origin** | **Specimens**<br>**/shipment** | **€/specimen** |
| --- | --- | --- |
| Indonesia | 765 | 2.6 |
| Philippines | 797 | 1.7 |
| Sri Lanka | 302 | 3.2 |
| United States | 333 | 5.6 |
| Kenya | 504 | 4.0 |
| Maldives | 597 | 2.0 |
| Singapore | 301 | 1.1 |
| Israel | 364 | 1.7 |
| Fiji | 801 | 1.1 |
| Dominican<br>Republic | 655 | 0.7 |

In total, 43,582 shipments with 25,503,345 specimens were imported into Europe with an average of 3,187,918 specimens a year (Figure 5). The annual number of imported specimens has decreased by 59.9% between 2014 to 2021 (Figure 5 and 6). Although the number of specimens from Indonesia has decreased between 2014 and 2021, their overall value increased (Figure 7). For Kenya, the number of specimens exported increased but not in the same proportion as their value (Figure 7). Thirty European countries imported marine ornamental fishes between 2014 and 2021 including Iceland, Norway, San Marino and Switzerland which are not part of the EU, along with the United Kingdom which left the EU the end of 2020 (Table 4).

**FIGURE 7.**
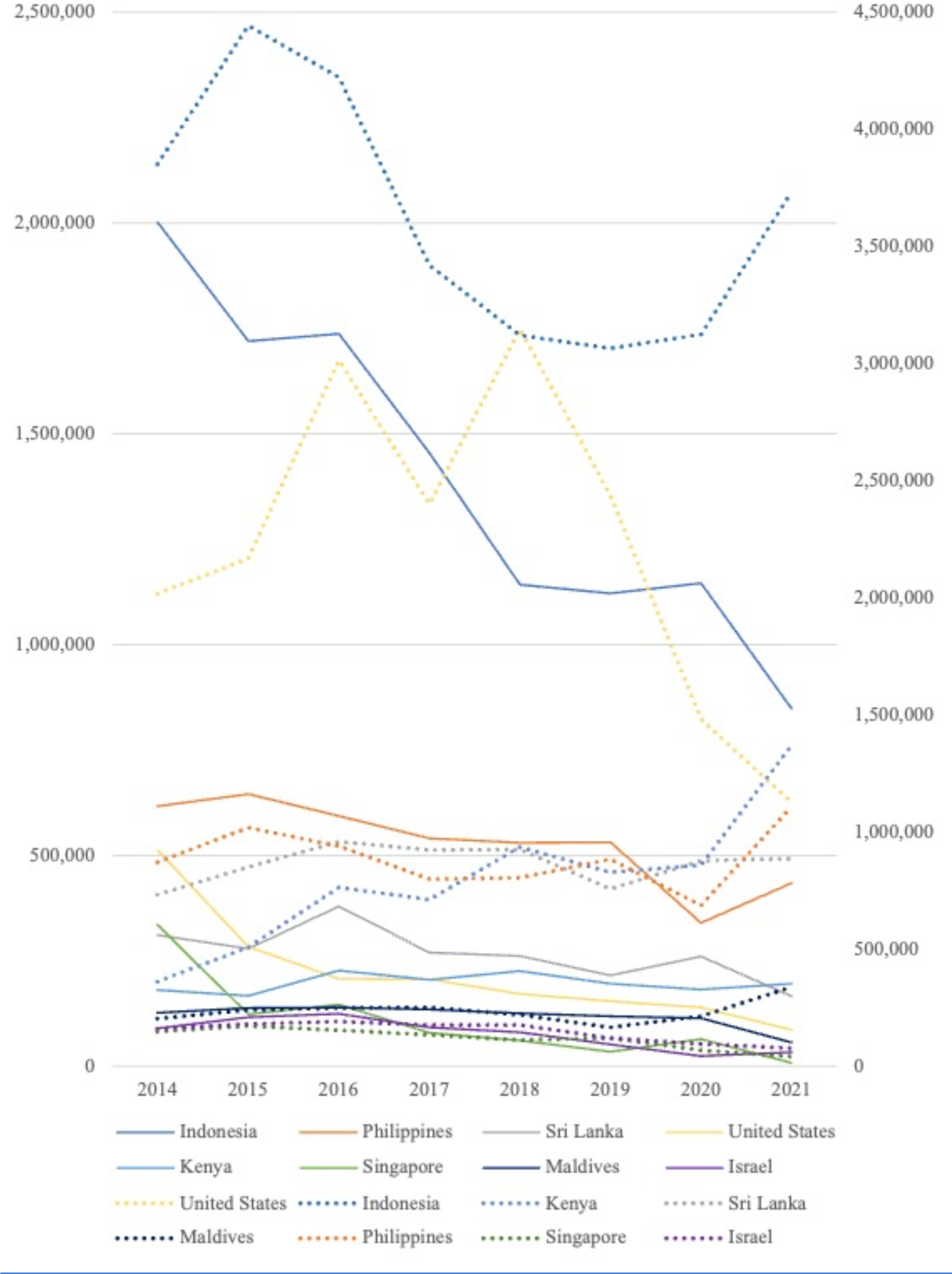
Number of specimens of marine ornamental fishes traded and value per year of the top eight exporting countries from 2014 to 2021 entering Europe according to TRACES data. Straight line = specimens, dotted line = value (Euro).

**TABLE 4.** Number of imported marine ornamental fishes per European country between 2014 and 2021 with total number of specimens over eight years.

| Country of origin | 2014 | 2015 | 2016 | 2017 | 2018 | 2019 | 2020 | 2021 | Total |
| --- | --- | --- | --- | --- | --- | --- | --- | --- | --- |
| United Kingdom | 1,414,494 | 1,034,680 | 978,506 | 844,667 | 791,320 | 734,017 | 677,486 | 5,589 | 6,480,759 |
| Netherlands | 1,049,798 | 864,511 | 828,314 | 567,534 | 434,157 | 397,051 | 410,299 | 555,299 | 5,106,963 |
| Germany | 583,889 | 482,637 | 660,114 | 551,266 | 404,108 | 432,017 | 411,755 | 416,326 | 3,942,112 |
| Italy | 431,839 | 379,421 | 373,391 | 320,154 | 328,957 | 295,946 | 226,673 | 258,509 | 2,614,890 |
| France | 487,863 | 377,900 | 305,278 | 261,525 | 240,181 | 257,648 | 249,002 | 195,783 | 2,375,180 |
| Spain | 237,842 | 117,045 | 131,727 | 81,042 | 129,689 | 102,110 | 49,573 | 46,768 | 895,796 |
| Poland | 93,191 | 78,044 | 87,300 | 94,473 | 85,277 | 96,609 | 119,399 | 106,882 | 761,175 |
| Denmark | 68,144 | 120,674 | 131,517 | 157,571 | 66,279 | 43,025 | 150,020 | 11,731 | 748,961 |
| Belgium | 99,186 | 62,984 | 76,810 | 70,276 | 59,691 | 48,769 | 67,871 | 39,246 | 524,833 |
| Sweden | 30,922 | 48,561 | 42,061 | 81,367 | 33,404 | 24,106 | 31,255 | 12,912 | 304,588 |
| Switzerland | 38,407 | 37,217 | 42,152 | 43,138 | 34,650 | 29,855 | 27,732 | 28,298 | 281,449 |
| Czech Republic | 22,187 | 27,462 | 26,633 | 21,817 | 41,022 | 31,291 | 60,351 | 34,426 | 265,189 |
| Portugal | 10,067 | 23,633 | 23,900 | 17,178 | 30,136 | 11,809 | 28,998 | 118,660 | 264,381 |
| Austria | 33,803 | 34,032 | 35,225 | 34,999 | 36,200 | 23,270 | 19,752 | 24,490 | 241,771 |
| Greece | 45,573 | 31,876 | 26,898 | 21,880 | 22,383 | 16,707 | 13,275 | 16,538 | 195,130 |
| Norway | 48,171 | 26,843 | 33,440 | 20,321 | 17,116 | 12,766 | 3,507 | 7,244 | 169,408 |
| Hungary | 20,442 | 7,065 | 6,834 | 9,797 | 9,670 | 9,795 | 11,049 | 7,044 | 81,696 |
| Luxembourg | 13,311 | 12,248 | 10,319 | 6,683 | 6,536 | 7,339 | 1,958 | NA | 58,394 |
| Romania | 1,650 | 1,344 | 1,613 | 6,124 | 19,121 | 13,882 | 6,774 | 523 | 51,031 |
| Cyprus | 6,576 | 6,068 | 4,735 | 5,566 | 4,102 | 13,926 | 4,138 | 4,339 | 49,450 |
| Malta | 4,992 | 10,130 | 5,530 | 5,476 | 3,835 | 9,943 | 1,786 | 1,839 | 43,531 |
| Bulgaria | 2,517 | 766 | 909 | 1,681 | 2,333 | 1,637 | 419 | 887 | 11,149 |
| Ireland | 838 | 190 | 826 | 175 | 397 | 281 | 2,313 | 6,005 | 11,025 |
| Other |  |  |  |  |  |  |  |  | 6,983 |
| Croatia | 481 | 879 | 519 | 199 | 958 | 1,584 | 1,072 | 1,270 | 6,962 |
| San Marino | NA | 1,801 | 2,392 | NA | NA | NA | NA | NA | 4,193 |
| Slovenia | 953 | 770 | 1,041 | 182 | 144 | NA | NA | NA | 3,090 |
| Iceland | NA | NA | NA | NA | 235 | 213 | 595 | 1,657 | 2,700 |
| Estonia | NA | NA | NA | NA | NA | 399 | NA | NA | 399 |
| Finland | NA | NA | NA | NA | NA | 96 | NA | NA | 96 |
| Slovakia | NA | 61 | NA | NA | NA | NA | NA | NA | 61 |

The EU country importing most marine ornamental fishes was the United Kingdom (6,480,759 specimens), except for 2021; it was followed by the Netherlands (5,106,963 specimens) that in 2021 imported more than the United Kingdom. The Netherlands was followed by Germany at 3,942,112 specimens. These three countries accounted for 60.9% of all imports of marine ornamental fishes into the EU between 2014 and 2021 (Table 4.). With the inclusion of Italy and France, these five countries alone imported a total of 80.5% of all marine ornamental fishes imported to the EU (Figure 8, Table 4).

**FIGURE 8.**
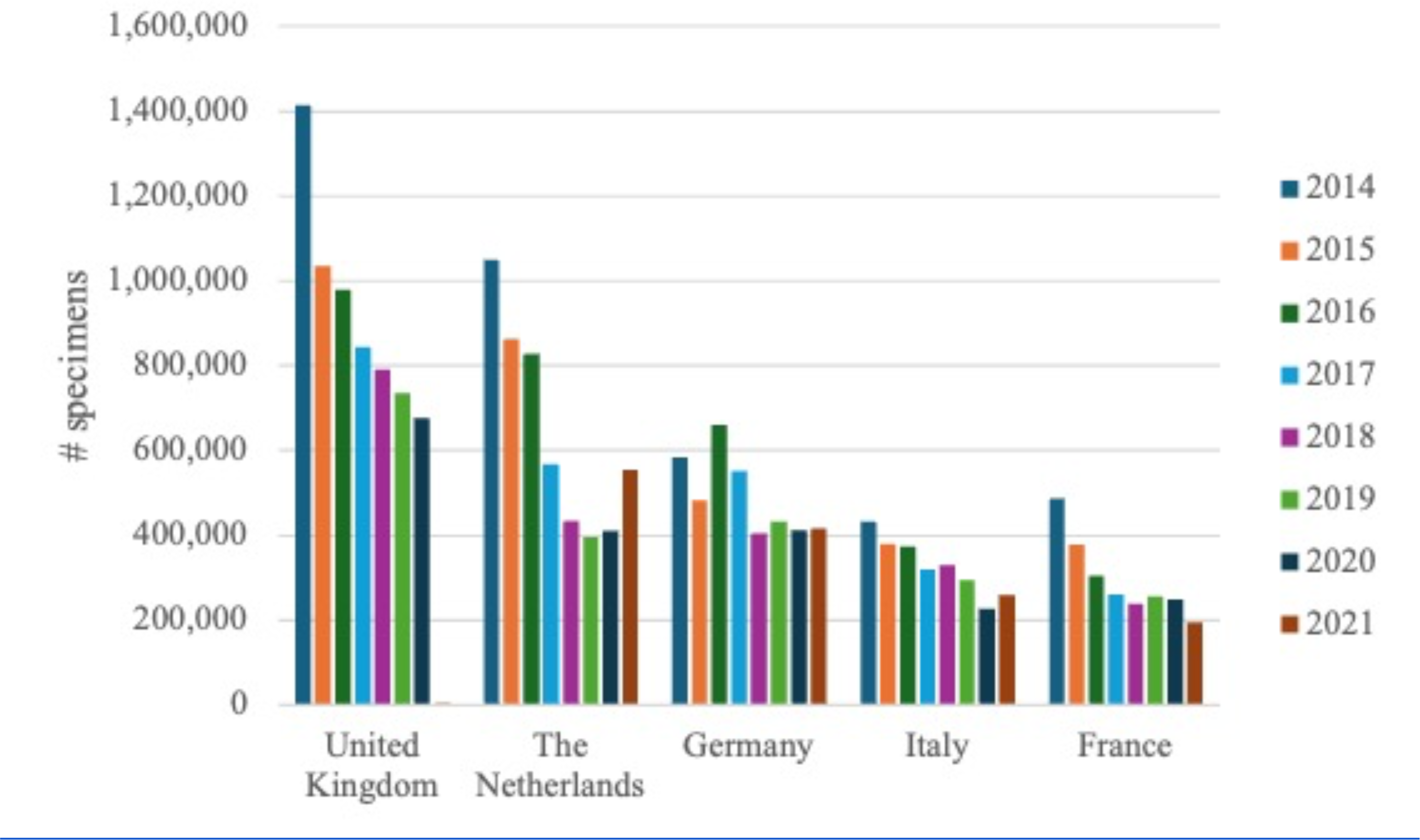
Number of specimens of marine ornamental fishes imported to the five top importing European countries by year from 2014 to 2021 according to TRACES data. The United Kingdom left the EU the end of 2020.

### Diversity of imported marine ornamental fishes

Between 2014 and 2021, fish species from 120 families were imported to Europe. The top 12 families accounted for 92.4% of all traded marine ornamental fishes in Europe number of specimens (Figure 9). Family Labridae featured the highest number of imported species (210), followed by the Pomacentridae (142), which was also the most traded family in number of specimens (Figure 9).

**FIGURE 9.**
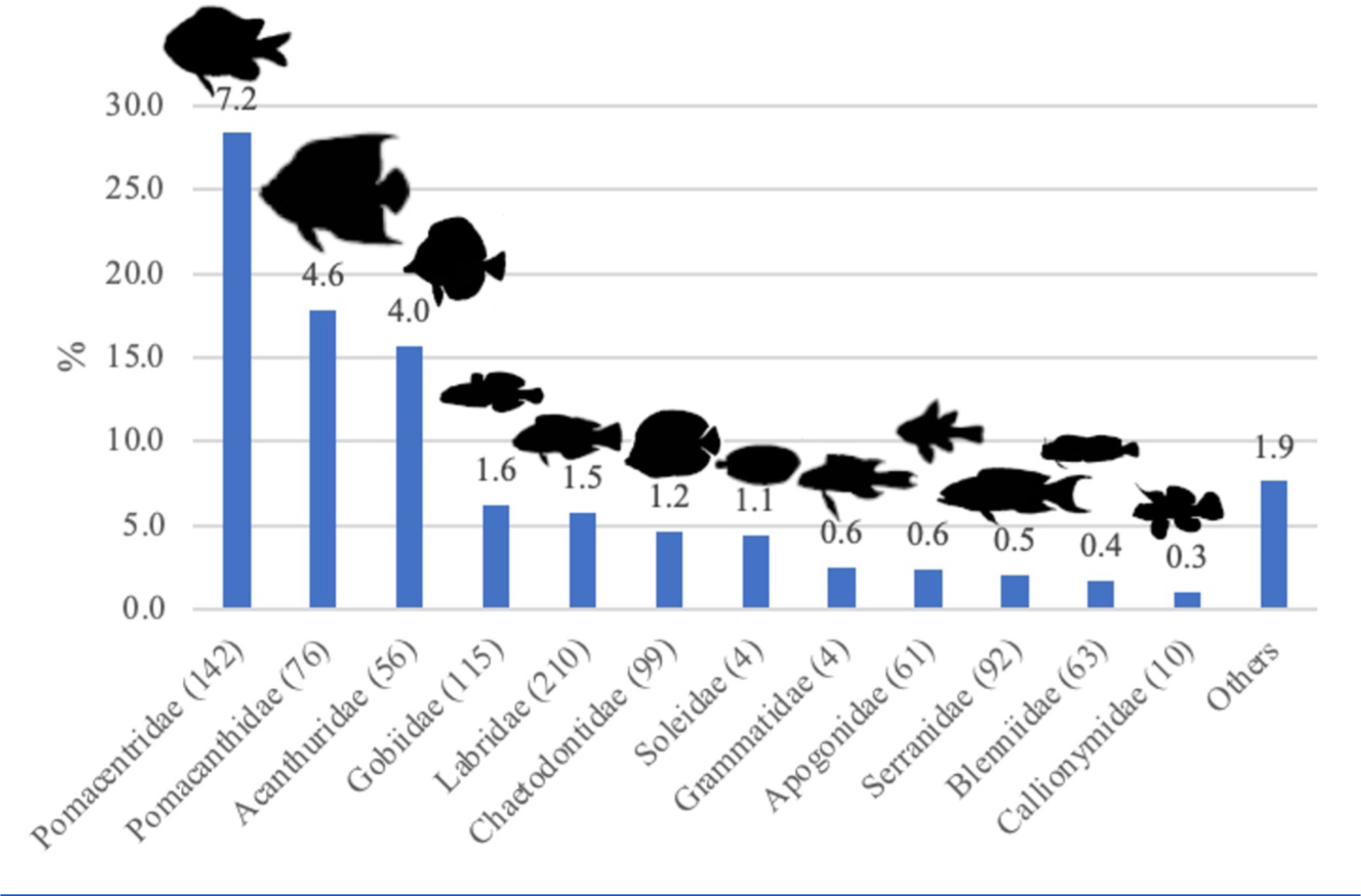
Number of specimens of marine ornamental fishes (% of total imported specimens) of the top twelve families traded into Europe between 2014 and 2021 according to TRACES data. The number of imported species per family is presented in parentheses, with the number on top of the bar representing millions of specimens.

Between 2014 and 2021, 1,452 species of marine ornamental fishes were imported to Europe. However, of the 25,503,345 specimens imported, only 17,770,326 specimens (69.7%) were registered at species level in the TRACES database (Table 1).

The blue green damselfish *Chromis viridis* was the most imported species, comprising 12.4% of the total number of imported marine ornamental fishes, followed by the clown anemonefish *Amphiprion ocellaris* with 10.0% and the bicolor angelfish *Centropyge bicolor* with 9.4% and of the total number of specimens (Table 5). The 20 most traded species accounted for 63.7% of the overall number of specimens imported into the EU between 2014 and 2021 where the species was known (Table 5, see supplementary material). Of the 20 most traded species, a total of 19 were listed as being of “least concern” by the IUCN Red List, with only the Banggai cardinalfish *Pterapogon kauderni*, ranked as the 9^th^ most imported species, being considered as “endangered” (Table 5). It is also worth highlighting that 14 of the 20 most traded species listed had been last evaluated ≥10 years ago (Table 5), three had an IUCN Red List population trend as “decreasing” and 7 “unknown” (Table 5).

**TABLE 5.**
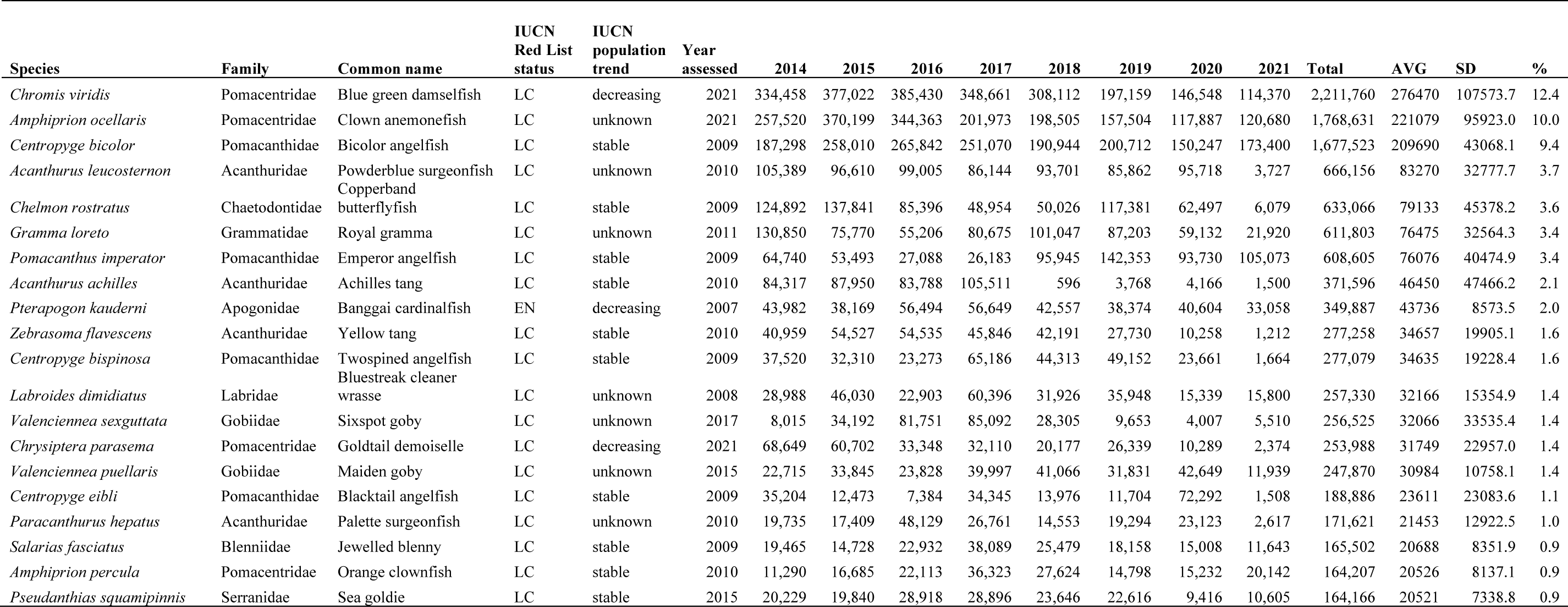
Top 20 species of marine ornamental fishes imported to Europe between 2014 and 2021 and their IUCN Red List conservation status (LC = “least concern”, EN = “endangered”), population trend as well as year assessed. AVG = average, SD = standard deviation, % 3 = percentage of traded specimens.

The IUCN Red List conservation status of all 1,452 species traded between 2014 and 2021 showed 1.3% to be “data deficient” or “not evaluated”, 95.5% as being “least concern”, 2% of species as “endangered” and only three species with 102 specimens being “critically endangered” (Table 6). The three “critically endangered” species imported were the scalloped hammerhead *Sphyrna lewini* (listed on CITES Appendix II) with 23 specimens being imported in 2015 from Philippines destined to France and 75 specimens in 2018 (2 from Kenya destined to France, 62 from Australia to the Netherland and 11 from Singapore to the Netherlands). Two Nassau groupers *Epinephelus striatus* were imported in 2016 from the Philippines destined for Italy. Sand tiger sharks *Carcharias taurus* were imported twice in 2020, both from the United States to the Netherlands (Table 6). No species traded were listed as “extinct in the wild” or “extinct”.

**TABLE 6.** IUCN Red List conservation status of all species of marine ornamental fishes and number of specimens per species imported into Europe between 2018 and 2021, as well as their % of the number of known marine ornamental fish species in Europe and worldwide according to FishBase. NE = “not evaluated”, DD = “data deficient”, LC = “least concern”, NT = “near threatened”, VU = “vulnerable”, EN = “endangered”, CR = “critically endangered”.

| **IUCN**<br>**Red List**<br>**status** | **2014** | **2015** | **2016** | **2017** | **2018** | **2019** | **2020** | **2021** | **Total** | **%** | **#**<br>**species** |
| --- | --- | --- | --- | --- | --- | --- | --- | --- | --- | --- | --- |
| NE | 17,671 | 26,354 | 16,538 | 12,713 | 11,412 | 15,027 | 8,092 | 12,341 | 120,148 | 0.7 | 63 |
| DD | 19,623 | 33,136 | 9,779 | 10,315 | 14,308 | 10,909 | 7,436 | 5,027 | 110,533 | 0.6 | 53 |
| LC | 2,754,733 | 2,697,184 | 2,643,394 | 2,399,028 | 2,121,468 | 1,936,373 | 1,578,146 | 911,066 | 17,041,392 | 95.5 | 1274 |
| NT | 5,704 | 4,181 | 3,114 | 2,465 | 595 | 1,537 | 683 | 300 | 18,579 | 0.1 | 18 |
| VU | 53,851 | 34,234 | 18,156 | 30,120 | 17,284 | 11,946 | 19,723 | 15,029 | 200,343 | 1.1 | 31 |
| EN | 44,000 | 38,195 | 56,526 | 56,686 | 42,570 | 38,421 | 49,807 | 33,088 | 359,293 | 2.0 | 10 |
| CR | NA | 23 | 2 | NA | 75 | NA | 2 | NA | 102 | 0.0 | 3 |

### Watchlists

The Watchlist alarm system gives a ranking of traded species (where species is known) from 2014 to 2021 based on number of trades specimens, IUCN Red List conservation status and vulnerability according to FishBase. The first 10 species are either CR or EN, with all species in trade already listed in CITES appendix II being present in the first 40 species of the Watchlist. Thirty-one species of pipefishes and seahorses (Syngnathidae) are on the Watchlist which are also listed on CITES appendix II (Table 7; supplementary material).

**TABLE 7.**
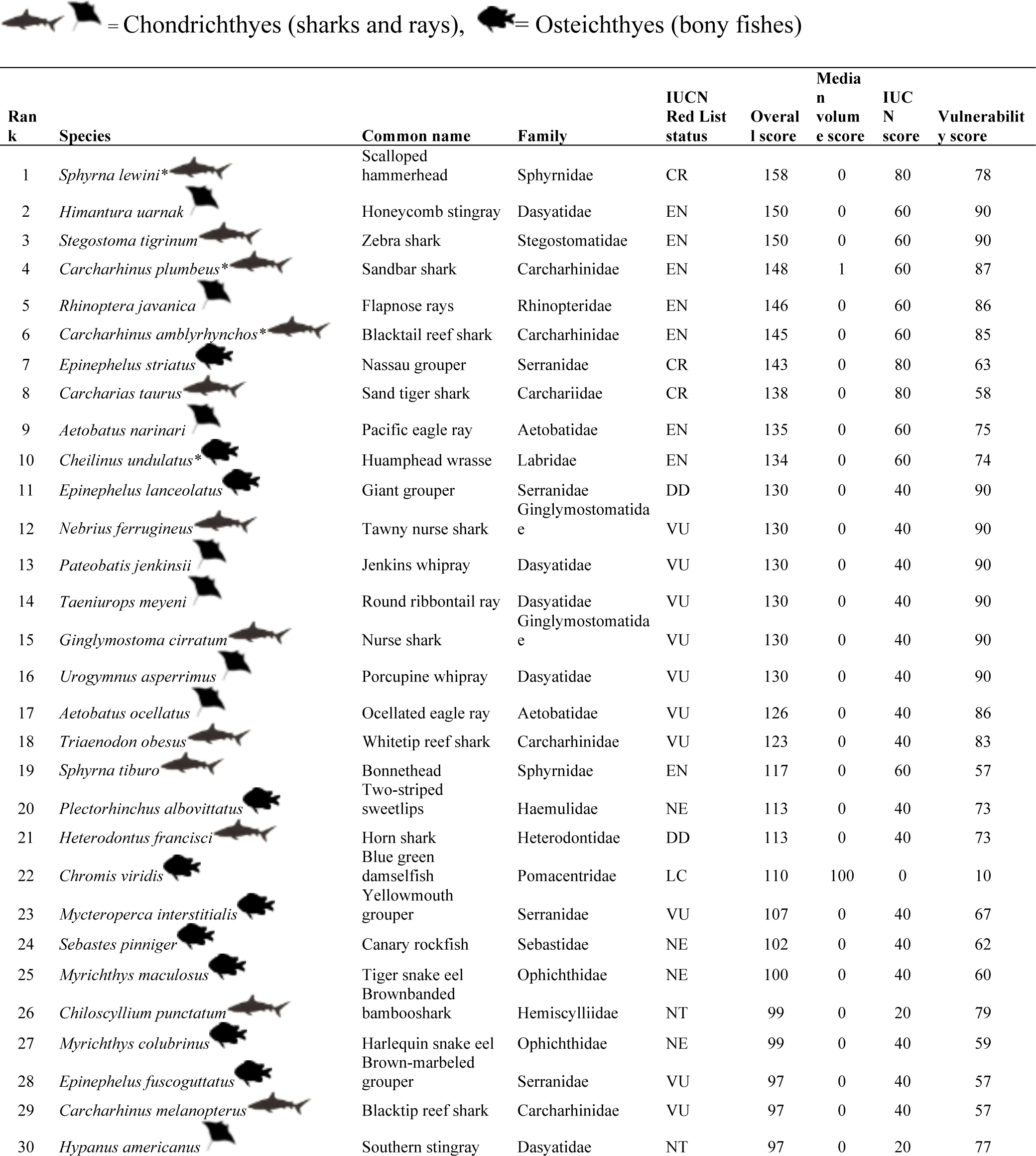
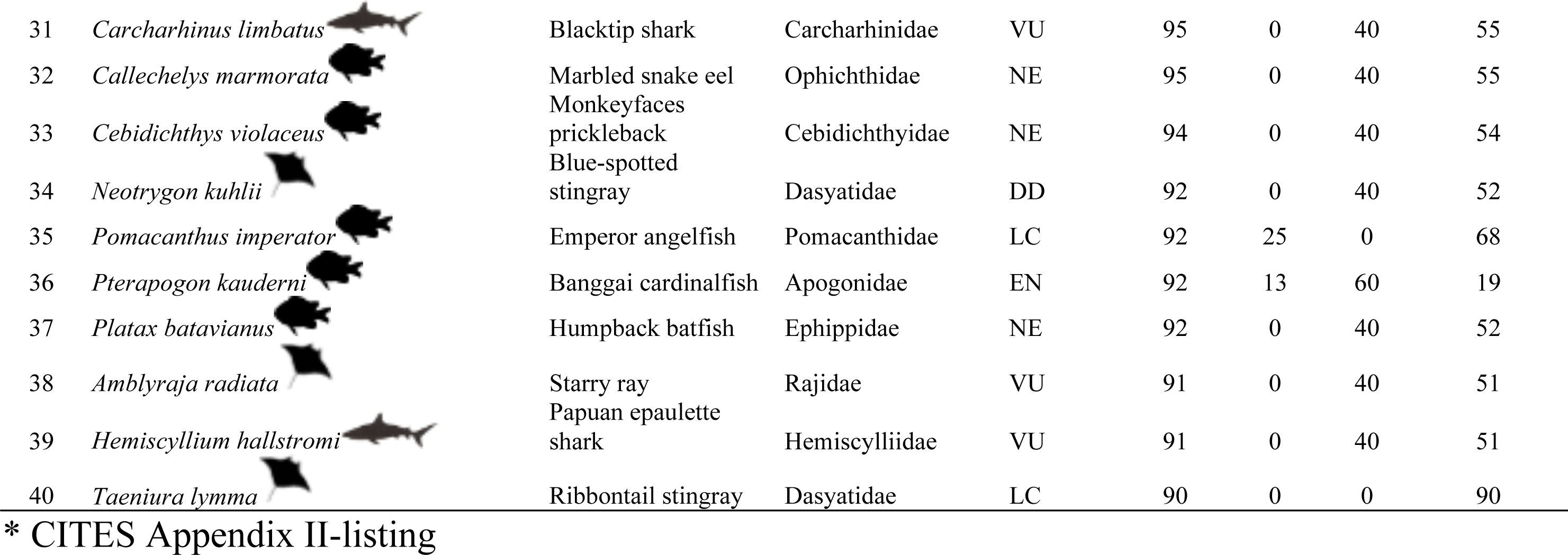
First 40 species with their IUCN Red List conservation status on the Watchlist ranked by the sum of three normalized parameters according to TRACES data from 2014 to 2021: the score of the median number of specimens traded, the score of the IUCN Red List conservation status (www.iucnredlist.org) and the score in vulnerability according to FishBase (http://www.fishbase.org) resulting in an overall score.

The most traded species, the blue green damselfish *Chromis viridis*, leads the ranking of the Watchlist+ which included the linear regression (slope of number of traded specimens over 8 years). It is followed by the Clown anemonefish *Amphiprion ocellaris* and the Copperband butterflyfish *Chelmon rostratus.* Unlike in the Watchlist, no cartilaginous appear on the Watchlist+, as they either been traded in very low numbers, or the number of species traded largely fluctuated over consecutive years thus making the linear regression not meaningful (Table 8).

**TABLE 8.** First 40 species with their IUCN Red List conservation status on the Watchlist+ ranked by the sum of four normalized parameters according to TRACES data from 2014 to 2021: the score of the median number of specimens traded, the score of the linear regression (slope) according to TRACES data, the score of the IUCN Red List conservation status (www.iucnredlist.org) and the score in vulnerability according to FishBase (http://www.fishbase.org) resulting in an overall score.

| Rank | Species | Common name | Family | IUCN Red List status | Overall score | Median volume score | Slope score | IUCN score | Vulnerability score |
| --- | --- | --- | --- | --- | --- | --- | --- | --- | --- |
| 1 | *Chromis viridis* | Blue green damselfish | Pomacentridae | LC | 210 | 100 | 100 | 0 | 10 |
| 2 | *Amphiprion ocellaris* | Clown anemonefish | Pomacentridae | LC | 157 | 62 | 84 | 0 | 10 |
| 3 | *Chelmon rostratus* | Copperband butterflyfish | Chaetodontidae | LC | 67 | 23 | 34 | 0 | 10 |
| 4 | *Acanthurus achilles* | Achilles tang | Acanthuridae | LC | 68 | 14 | 41 | 0 | 14 |
| 5 | *Zebrasoma flavescens* | Yellow tang | Acanthuridae | LC | 88 | 13 | 18 | 0 | 57 |
| 6 | *Chrysiptera parasema* | Goldtail demoiselle | Pomacentridae | LC | 42 | 9 | 23 | 0 | 10 |
| 7 | *Chrysiptera cyanea* | Sapphire devil | Pomacentridae | LC | 28 | 6 | 13 | 0 | 10 |
| 8 | *Nemateleotris magnifica* | Fire goby | Gobiidae | LC | 21 | 5 | 6 | 0 | 10 |
| 9 | *Centropyge loriculus* | Flame angel | Pomacanthidae | LC | 25 | 5 | 10 | 0 | 10 |
| 10 | *Zebrasoma velifer* | Sailfin tang | Acanthuridae | LC | 59 | 4 | 19 | 0 | 37 |
| 11 | *Centropyge acanthops* | Orangeback angelfish | Pomacanthidae | LC | 28 | 4 | 14 | 0 | 10 |
| 12 | *Acanthurus japonicus* | Japan surgeonfish | Acanthuridae | LC | 27 | 3 | 13 | 0 | 11 |
| 13 | *Pomacanthus annularis* | Bluering angelfish | Pomacanthidae | LC | 44 | 3 | 6 | 0 | 35 |
| 14 | *Naso lituratus* | Orangespine unicornfish | Acanthuridae | LC | 39 | 2 | 3 | 0 | 34 |
| 15 | *Pomacentrus alleni* | Adaman damsel | Pomacentridae | LC | 16 | 2 | 4 | 0 | 10 |
| 16 | *Pseudochromis fridmani* | Orchid dottyback | Pseudochromidae | LC | 15 | 2 | 3 | 0 | 10 |
| 17 | *Centropyge tibicen* | Keyhole angelfish | Pomacanthidae | LC | 17 | 2 | 5 | 0 | 10 |
| 18 | *Centropyge potteri* | Russet angelfish | Pomacanthidae | LC | 17 | 2 | 5 | 0 | 10 |
| 19 | *Halichoeres chrysus* | Canary wrasse | Labridae | LC | 14 | 2 | 2 | 0 | 10 |
| 20 | *Dascyllus aruanus* | Whitetail dascyllus | Pomacentridae | LC | 31 | 1 | 3 | 0 | 26 |
| 21 | *Amphiprion polymnus* | Saddleback clownfish | Pomacentridae | LC | 16 | 1 | 4 | 0 | 10 |
| 22 | *Dascyllus trimaculatus* | Treespot dascyllus | Pomacentridae | LC | 16 | 1 | 5 | 0 | 10 |
| 23 | *Ecsenius bicolor* | Bicolor blenny | Blenniidae | LC | 13 | 1 | 2 | 0 | 10 |
| 24 | *Chaetodon auriga* | Threadfin butterflyfish | Chaetodontidae | LC | 32 | 1 | 7 | 0 | 23 |
| 25 | *Dascyllus melanurus* | Blacktail humbug | Pomacentridae | LC | 15 | 1 | 4 | 0 | 10 |
| 26 | *Pomacentrus coelestis* | Neon damselfish | Pomacentridae | LC | 13 | 1 | 2 | 0 | 10 |
| 27 | *Neosynchiropus ocellatus* | Ocellated dragonet | Callionymidae | N.E. | 54 | 1 | 2 | 40 | 10 |
| 28 | *Synchiropus picturatus* | Picturesque dragonet | Callionymidae | LC | 13 | 1 | 2 | 0 | 10 |
| 29 | *Heniochus acuminatus* | Pennant coralfish | Chaetodontidae | LC | 20 | 1 | 4 | 0 | 15 |
| 30 | *Acanthurus olivaceus* | Orangespot surgeonfish | Acanthuridae | LC | 20 | 1 | 1 | 0 | 18 |
| 31 | *Amblygobius phalaena* | Whitebarred goby | Gobiidae | LC | 13 | 1 | 2 | 0 | 10 |
| 32 | *Canthigaster valentini* | Valentin's sharpnose puffer | Tetraodontidae | LC | 12 | 1 | 1 | 0 | 10 |
| 33 | *Haemulon flavolineatum* | Frenchg grunt | Haemulidae | LC | 44 | 1 | 12 | 0 | 32 |
| 34 | *Pterois volitans* | Red lionfish | Scorpaenidae | LC | 37 | 1 | 2 | 0 | 34 |
| 35 | *Odonus niger* | Red-toothed triggerfish | Balistidae | LC | 42 | 1 | 1 | 0 | 40 |
| 36 | *Holacanthus ciliaris* | Queen angelfish | Pomacanthidae | LC | 38 | 0 | 2 | 0 | 35 |
| 37 | *Centropyge flavissima* | Lemonpeel angelfish | Pomacanthidae | LC | 12 | 0 | 2 | 0 | 10 |
| 38 | *Chrysiptera rollandi* | Rolland's demoiselle | Pomacentridae | LC | 11 | 0 | 1 | 0 | 10 |
| 39 | *Amphiprion allardi* | Twobar anemonefish | Pomacentridae | LC | 11 | 0 | 0 | 0 | 10 |
| 40 | *Synchiropus morrisoni* | Morrison's dragonet | Callionymidae | LC | 11 | 0 | 0 | 0 | 10 |

## Discussion

The EU is a major global economic market and a leading importer of marine ornamental fishes. In this way, it is legitimate to say that the EU plays a crucial role in ensuring the sustainability of this trade. Indeed, the EU has taken significant steps in this regard, including the “Revised EU Action Plan to End Wildlife Trafficking” (EU 2022b) and the “European Animals Health Law”, which addresses disease transmission risks in wild animals (EU 2016). Also, the EU together with the US and Switzerland asked for this trade to be scrutinized at the 18^th^ conference of the Parties to CITES in August 2019. In this sense, it is a logical continuation of these actions to adapt TRACES to properly monitor the trade in ornamental marine fish. The EU Parliament’s resolution of 5 October 2022 stressed the importance of addressing the trade of marine ornamental fishes. It urged the Commission to modify the European TRACES database, ensuring accurate and publicly accessible information on species, specimen numbers, and trade origins (EU 2022b).

### The European Trade Control and Expert System TRACES

Although TRACES is not a monitoring tool to specifically target wildlife, it provides valuable data for estimating the number of specimens and species diversity in the trade of marine ornamental fishes imported to Europe (TRACES 2023). TRACES, implemented in the EU 2004, became applicable for monitoring the marine ornamental fish trade in 2014 (TRACES 2023). All marine ornamental fishes imported to Europe arrive via air freight and undergo customs clearance and veterinary inspection upon arrival. Documents accompanying the shipments provide species-level information which may be more specific in taxonomic detail from the electronically filled out TRACES information. This more detailed information can be easily inputted into TRACES without imposing excessive workload on users (Gillett et al. 2020). In a survey conducted in 2008, industry representatives expressed support for trade monitoring through veterinary controls, as the forms used already request species-level information and are routinely completed (UNEP-WCMC 2008). Regrettably, nothing came off this survey.

Moreover, TRACES allows for electronic data import, alignment with FAIR data principles (data which are findable, accessible, interoperable, and reusable) and accurate scientific identification of fish species. Also, it could potentially be used for monitoring other vertebrate taxa as well, and its fine tuning for this purpose could easily be tested by using marine ornamental fishes as a case study. However, an update in 2019 resulted in less accurate data collection for marine ornamental fishes. A significant portion of specimens imported between 2018 and 2021 were incorrectly registered in TRACES as marine ornamental fishes when these were actually freshwater fishes, invertebrates, or even amphibians. Previous analysis from 2014 to 2017 had a minimal number of shipments with no information, which were excluded from the study (Biondo and Burki 2019). Until 2019, most inaccuracies only occurred at a family level, but at present genera, families or even orders can be registered in this platform which surprisingly makes the updated version of TRACES less accurate and more challenging to analyze. Nonetheless, this updated version is also easier to handle, as data files can be imported rather than having to type information by hand.

It is also worth noting that monitoring systems outside of Europe are often more inaccurate than TRACES (Rhyne et al. 2012; Trujillo-González and Militz 2019). For instance, the United States relies on the Law Enforcement Management Information System (LEMIS), which lacks taxonomic detail and reports most of the trade as generic categories (less than 0.2% or fewer than 22,000 individuals/year at species or genus level) (UNEP-WCMC 2022). Australia’s monitoring system also has very limited taxonomic resolution, leaving uncertainties about the true nature of their imports (Trujillo-González and Militz 2019). Extracting information based on weight (kg/year) from databases such as UN Comtrade (UN 2023) has been suggested as a proxy to monitor this trade, but this approach includes using the weight of water on which fishes are shipped, thus making this figure highly unreliable for determining the number of marine ornamental fishes being imported/exported (UNEP-WCMC 2022). The lack of species-specific information not only hampers scientific analysis but also poses a biosecurity risk, which the EU aims to address through its Animal Health Law (EU 2016). Australia recognizes the risks associated with the trade of ornamental fishes and has implemented strict import biosecurity measures to control diseases (Hood et al., 2019; Johan and Zainathan 2020).

### Import values

According to European imports and total demand, marine ornamental fishes are becoming more expensive. The global trade of marine ornamental fishes has always been much more valuable than that of food fishes (Watson et al. 2023). In the 1980s, marine aquarium fishes were priced at US$ 750/kg, while marine food fishes were priced at US$ 9/kg (Holcombe et al. 2022). Presently, marine ornamental fishes are valued at a minimum of US$ 1,000/kg compared to US$ 13/kg for marine food fish (Leingang 2021). Coral reefs contribute US$ 2.7 trillion annually in goods and services, including US$ 36 billion from coral reef tourism (Dey 2016; Souter et al. 2021). The industry of aquatic organisms for home and public aquariums, along with the equipment required to display these organisms, is estimated to be a multi-billion-dollar industry (Stevens et al. 2017). In the 1980s, the global ornamental fish industry with associated equipment and accessories was valued at US$ 7.2 billion (Andrews 1990), which increased to US$ 20-30 billion by 1997 (Biondo and Burki 2019; Dey 2016; Penning et al. 2009; Raghavan et al. 2013; Surtida 1999; Teletchea 2016). By 2004, the estimated value ranged from US$ 800 million to US$ 30 billion annually (Biondo and Burki 2019; Ploeg 2007; Saxby et al. 2010; Stevens et al. 2017; Whittington and Chong 2007). Currently, the import value of marine ornamental fishes solely shipped to and within Europe is € 24.7 million while total demand is about € 37.6 million. On the other hand, extra EU-imports diminished from € 13 million to € 9.5 million between 2014 and 2021. The latter figure is similar to extra-EU import estimates previously reported for the period 2000-2011 (Leal et al. 2015). This figure is likely to be more comparable to the import levels of regions where intra-regional trade is not accounted for in the data (especially the case of China in this analysis). However, and although the number of specimens recorded by TRACES has decreased, the value of intra-EU imports into the EU as well as the value of total demand remain steady. The long-term trend of the EU final demand with regards to value does not appear to be diminishing, and therefore the reduction in the number of specimens may be a result of the market moving into “higher-value market niches” or the result of a rigid supply being unable to meet demand (as the increase in average prices suggests). The sharp increase in European average prices in 2021 accelerated the increasing trend in prices, most likely as a result of the Covid-19 pandemic. The apparent change may also be due to the declining quality of the data. Increasing import values in non-EU countries, mainly China, the US, and the Middle East suggest that the international demand for marine ornamental fishes will continue to expand, pushing prices up and increasing the pressure on local fish populations. It is therefore crucial to recognize that live fishes are just one part of a larger commercial system that includes food, supplements, and a wide range of accessories and equipment. Unequivocally, the value of this system is much greater than that of live fishes alone (Biondo and Burki 2020). The extensive economic impact of this integrated trade, supported by a complex network of suppliers, traders, retailers, and consumers, puts significant pressure on marine fishes used as ornamentals. This emphasizes the need for improved trade reporting and monitoring systems. Overall, fewer specimens per year were imported to Europe from 2014 to 2021 at an almost constant decreasing rate. As the import value stayed the same over the eight years, each fish specimen got more expensive. A reason for a product to get more expensive is if the supply is becoming scarcer, often due to a population decrease when it comes to live specimens. Another explanation for an increase in price would be due to a higher demand driven by other countries, such as China, and supply being unable to keep up.

### Assessment of environmental consequences

Habitat loss is the greatest threat to biodiversity (Pinho et al. 2020) and coral reefs are among the most threatened marine ecosystems due to anthropogenic interferences (Halpern et al., 2007, IPCC 2018; Souter et al., 2020). The Intergovernmental Science-Policy Platform on Biodiversity and Ecosystem Services (IPBES) has identified overexploitation, including trade, as the second leading cause of extinction for nearly one million species (IPBES 2019). Estimates suggest that global trade involves 15 to 30 million coral reef fishes annually (Biondo and Burki 2020), with potential figures reaching as high as 150 million (Stevens et al. 2017). Marine ornamental fishes have yet to be included in CITES appendix I (trade ban), with only seahorses, humphead wrasse, and clarion angelfish as well as a few sharks and rays monitored under CITES Appendix II (monitored trade).

Almost all marine ornamental fishes traded worldwide are still sourced from the wild, mostly from coral reefs, as they have complex life cycles that are difficult to replicate in breeding facilities or aquariums (Biondo 2018, 201a; Biondo and Burki 2019, 2020; Pouil et al. 2020; Rhyne et al. 2012, 2017). As of 2018, only 24 species were bred in commercial numbers in captivity, while 338 had already been bred at what can be considered a more academic level; indeed, the number of species bred in captivity has remained fairly constant since 2012 (Pouil et al. 2020). Keeping marine ornamental fishes is expensive and demanding (OATA 2014). Furthermore, a survey of over 3,000 marine ornamental fish keepers showed that more than 70% did not intend to try to breed fishes (Pountney 2023).

### Origin, destination, diversity, and conservation

Overall, from 2014 to 2021, 61 countries exported marine ornamental fishes to Europe which represent an increase of 18% from 2014-2017 (Biondo and Burki 2019). Some small island countries, such as Palau, São Tomé and Príncipe, and Tonga did not appear to export to Europe the last few years of this analysis, despite having rare and highly sought-after species for the marine aquarium trade (Gillett et al. 2020). Singapore still plays a significant role as a transit hub for this trade, and it remains unclear where species labeled as originating from Singapore were actually collected in that country. It is important to note that China’s domestic wildlife trade is significant, emphasizing the need for a global monitoring. The Netherlands surpassed the UK as the leading importer in 2021 after the latter exiting the EU (an event popularly known as Brexit).

Species diversity (number) of imported marine ornamental species reach a total of 1,452 species over the whole period. Unfortunately, for one third of all specimens traded it was not possible to identify them to species level. Strangely, a CITES document analyzing TRACES data from 2018 to 2021 found only 33 species in TRACES, (UNEP-WCMC 2022), with some species using outdated names no longer accepted by WoRMS (e.g., *Abalistes stellaris* instead of *A. stellatus*) (W. Appeltans, 2023).

The total number of families imported over the whole period was 120 families which is higher than the number of 86 traded families since the last study (Biondo and Burki 2019). This rise may suggest that the most commonly available species in the trade may no longer be so abundant in the wild due to the dire state of the marine ornamental fishes’ habitats - tropical coral reefs (IPCC 2018; Souter et al. 2020; Hoegh-Guldberg 2019).

The number of the most traded species, the blue green damselfish *Chromis viridis*, has diminished significantly by 70%. The species was last evaluated in 2021 and is listed as “least concern” but the wild population is decreasing (Allen et al. 2022a). Also, the second most traded species, the clown anemonefish *Amphirion ocellaris* shows a decrease by almost 70% in eight years. This species was also last evaluated in 2021 and is of “least concern” according to the IUCN Red List, although its population status is unknown (Allen et al 2022b). These two species account for almost a quarter of the whole trade in Europe and explain the decreasing trend reported in this study. While *A. ocellaris* is increasingly bred in captivity in importing countries, and this may somehow explain why less specimens are imported into the EU, this is certainly bot the case of *C. viridis* (Pouil et al. 2020). The third most traded species is the bicolor angelfish *Centropyge bicolor* which, according to the IUCN Red List, has a stable population but was last evaluated in 2009. The trend recorded in the present study may be similar to that already recorded for food fishes in the past, where an increased diversity on species number occurred due to the scarcity of established food fish fisheries in commercial amounts (Hughes et al. 2022). Another potential explanation for this trend is that the marine aquarium trade regularly introduces new fish species to meet the demand for unique and novel organisms sought by hobbyists, an approach that aims to keep the price of “rare” species in the high-end range (Rhyne et al. 2014).

Our study allowed to confirm that “critically endangered” fish species were imported. For instance, in 2015 France imported 23 scalloped hammerhead *Sphyrna lewini* (listed on CITES Appendix II) from the Philippines, in 2016 Italy imported two “critically endangered” Nassau groupers *Epinephelus striatus* from the Philippines, and in 2018 75 specimens of scalloped hammerheads *Sphyrna lewini* were also imported: Two from Kenya destined to France, 62 from Australia destined to the Netherland and 11 more from Singapore also destined to the Netherlands. Three hammerhead sharks, recorded only at family level, were imported to Italy from Sri Lanka. Sand tiger sharks *Carcharias taurus* were imported twice in 2020, both from the United States to the Netherlands reflecting the presence of a major shark wholesaler in the country equipped with facilities to hold such large and active animals. Notably, eleven scalloped hammerheads had Singapore as their country of origin, highlighting Singapore’s role as a transit hub in the marine aquarium trade. This increase in live-shark trade is concerning, as nearly two-thirds of shark and ray species associated with coral reefs are at risk of extinction (Sherman et al. 2023) due the ongoing decline of shark species caused by commercial fishing and by-catch, despite all management efforts to revert this trend (Burgess et al. 2022).

Of the 1,452 marine ornamental fish species traded between 2014 and 2021, 1.3% had a conservation status of “data deficient” or “not evaluated. This is a stark decrease compared to 33.63% of the 1,334 species traded (with known species level) between 2014 and 2017 (Biondo and Burki 2019). While it is positive that fewer species require evaluation, the IUCN Red List advises not to assume that these categories indicate non-threatened status. Until assessed, it is recommended to give the same attention to data deficient species as to already recognized threatened species (IUCN 2023). Furthermore, for many species of marine ornamental fishes the IUCN Red List evaluation is outdated by over ten years for, leaving their current conservation status unclear. It is possible that “data deficient” species may be more threatened than initially perceived (Borgelt et al. 2022). Research indicates that a third of species listed as “not threatened” are experiencing a decline. Moreover, for nearly 75% of the 25,000 analyzed fish species, the population trend is unknown due to data limitations, which poses a significant challenge for understanding their status (Finn et al. 2023).

### Watchlist and Watchlist+ alarm system

The Watchlist considers the number of traded specimens, species vulnerability according to Fishbase, and IUCN Red List conservation status and gives a ranking using the overall scores, while the Watchlist+ further takes into account a linear regression of the number of specimens being annually traded. Both can be used as an indicator of potential negative impacts promoted by the international trade of marine ornamental species, with the Watchlist+ having more robust statistics regarding the trend of specimens traded. Both watchlists can serve as an alarm system, a starting point to identify species that require closer analysis or observation, whether due to the high number of specimens being traded or their ecological vulnerability that may lead to a possible population decline. Moreover, it also provides insights into which species could receive a better precautionary monitoring through CITES. Unfortunately, about a third of all imported marine ornamental fish specimens lack species identification and therefore could not be taken into consideration to enhance the magnitude of this watchlist approach.

In the Watchlist the blue green damselfish *Chromis viridis* ranked 22^nd^ representing the first Osteichthyes in the Watchlist but 1^st^ in the rankings in the Watchlist+ due to its large numbers in trade and the strongest decline recorded in the number of traded specimens. The blue-green damselfishes, initially considered a complex of two species (*C. viridis* and *Chromis atripectoralis*), were differentiated primarily based on the coloration of the pectoral fin base (Froukh and Kochzius 2008). Interestingly, *C. atripectoralis* was mainly exported from the Philippines, while imports of *C. viridis* listed the Caribbean (e.g., Cuba) as the country of origin. However, the Caribbean is not a natural habitat for this Indo-Pacific species. Both species leading the Watchlist+ show a pronounced decline in the number of traded specimens and definitely qualify for an in-depth analysis on the reasons why such trend was displayed. As the linear regression was not meaningful, the Banggai cardinalfish *Pterapogon kauderni*, ranked 36^th^ on the Watchlist,contrast to the Watchlist, no cartilaginous fishes, even if already CITES-listed on appendix II, made it to the Watchlist+, as they have been either traded in very low numbers or the number of species traded largely fluctuated over consecutive years. Similarly, the humphead wrasse *Cheilinus undularus*, which is also on CITES appendix II, also did not make it to the Watchlist+ due to the reduced numbers in trade, most likely due to its large size at adulthood, thus only being viable for display on public aquariums; nonetheless, as over half a million wrasses (Labridae) were not registered at species level, the accuracy of this finding must be put into perspective, reinforcing the need for a more accurate monitoring system.

### Advantages of TRACES and its adaptation to monitor wildlife

The trade of marine ornamental fishes can pose biosecurity risks, potentially leading to the unintentional spread of pathogens such as viruses (Hood et al. 2019; Johan and Zainathan 2020). Additionally, there is a hazard of introducing exotic species that may become invasive, exemplified by the red lionfish *Pterois volitans*. Despite their negative impacts, such as predation on smaller fishes and the establishment of large populations, lionfish specimens are still imported into the US (Lyons et al. 2019). The EU should take heed of this lesson and prohibit lionfish imports, as *Pterois* species are already present at the Mediterranean Sea (Kletou et al. 2016). While the import of the devil firefish *Pterois miles* was banned in 2015, Europe has imported over 300 specimens since 2016 (Kleitou et al. 2021), highlighting the need to enhance TRACES for a more comprehensive information at species-level.

The absence of species-level information in the long-standing marine ornamental fish trade has significant conservation implications, as highlighted in previous studies (Biondo, 2017 2018; Biondo and Burki 2019, 2020; UNEP-WCMC 2022; Wabnitz et al. 2003).

While TRACES is not specifically designed for monitoring trade in wildlife, including marine ornamental fishes, it already possesses the necessary features for such purposes, and the pet trade industry has already endorsed its use (UNEP-WCMC 2008). TRACES can be easily adapted to collect accurate data independently of CITES decisions, as traders are already required to submit information electronically and as an EU stakeholder survey suggests, they would be willing to do so (EU 2008). By 2014, TRACES recorded marine ornamental fishes with its specific “Harmonized System code 03011900 for Live ornamental fish (excluding freshwater)” category. Until 2019 it was also possible to list marine ornamental fishes at family level, which made data blurry. Since 2019, TRACES has become even more blurry, with almost 70% of specimens listed under the marine ornamental fish code being freshwater fishes, invertebrates, or amphibians. On the positive side, TRACES now allows the import of Excel tables, but unfortunately the database now also allows the import of information at a higher taxonomic level, such as orders.

TRACES should be fine-tuned so it would: a) only allow marine ornamental fishes under code “HS 03011900”, i.e. coral reef fishes according to the World Register of Marine Species (WoRMS; www.marinespecies.org); b) only accepting scientific names; c) making it mandatory to specify if specimens being traded were sourced from the wild by specifying geographic origin (region and country of capture) or if they were captive bred by providing the address of the breeding facility.

Moreover, a fine tunned TRACES may also contribute to help clarifying the potential confounding effects on the trade of marine ornamental fish promoted by both “Brexit” and the COVID-19 pandemics. The role to be played by the UK on this trade remains to be clarified in upcoming years, as well as confirming that the Netherlands has become the new “EU hub” for imports of marine ornamental fish. Concerning the UK, it is most likely that this country will continue to use TRACES (as other EU third parties already do, such as Norway or Switzerland) to record all its imports concerning live animals, as well as feed, plants and other goods, rather than establishing its own monitoring system. As such a follow up study to the present one covering data spanning until 2025 (or so), once these are made available by authorities, after proper data curation, will allow to uncover any potential bias promoted by the Brexit one the data presented in this study. The same rationale can also be applied to the COVID-19 pandemics, as only by confirming the trends reported for 2020 and 2021 in subsequent will it be possible to clarify if and how biased our data may have been from this unique event. Overall, a refined TRACES will be a valuable tool to accurately report reliable data on the trade of wildlife in general and live marine ornamental fish in particular in the EU.

## Supporting information

Watchlist+

Watchlist

Species specimens

## Acknowledgement

We are thankful to the EU for providing the raw data from TRACES. and to Rebekka Gammenthaler as well as Keith Lindsay for proofreading the manuscript.

## Supplementary material

Species traded and number of specimens Watchlist and Watchlist+

## Footnotes

1 Converted into €, total import values obtained from Comtrade are –5.9% lower (on average, 2014-2021) than those reported by the EuroStat for the same product classification. Part of the variation is due to movements in the exchange rate.

